# Single Object Profiles Regression Analysis (SOPRA): A novel method for analyzing high content cell-based screens

**DOI:** 10.1101/2021.09.24.461074

**Authors:** Rajendra Kumar Gurumurthy, Klaus-Peter Pleissner, Cindrilla Chumduri, Thomas F. Meyer, André P. Mäurer

## Abstract

**Motivation:** High content screening (HCS) experiments generate complex data from multiple object features for each cell within a treated population. Usually these data are analyzed by using population-averaged values of the features of interest, increasing the amount of false positives and the need for intensive follow-up validation. Therefore, there is a strong need for novel approaches with reproducible hit prediction by identifying significantly altered cell populations.

**Results:** Here we describe SOPRA, a workflow for analyzing image-based HCS data based on regression analysis of non-averaged object features from cell populations, which can be run on hundreds of samples using different cell features. Following plate-wise normalization the values are counted within predetermined binning intervals, generating unique frequency distribution profiles (histograms) for each population, which are then normalized to control populations. Statistically significant differences are identified using a regression model approach. Significantly changed profiles can be used to generate a heatmap from which altered cell populations with similar phenotypes are identified, enabling detection of siRNAs and compounds with the same ‘on-target’ profile, reducing the number of false positive hits. A screen for cell cycle progression was used to validate the workflow, which identified statistically significant changes induced by siRNA-mediated gene perturbations and chemical inhibitors of different cell cycle stages.

## Background

The availability of robotic liquid handling combined with automated fluorescence microscopy and high-performance image computing has enabled rapid advances in the development of high-throughput screening. Numerous studies have demonstrated the power of high-throughput image-based assays for characterizing drug effects (Perlman, et al., 2004), identifying active small molecules (Tanaka, et al., 2005) and classifying sub-cellular protein localization (Boland and Murphy, 2001; Conrad, et al., 2004), including genome-wide siRNA-mediated loss-of-function screens (Neumann, et al., 2006) or gene deletion (Ohya, et al., 2005) libraries. For each single cell within a cellular sample population, it is possible to achieve quantitative measurements of phenotypes such as expression level and localization of proteins, post-translational modifications and even cellular or sub-cellular morphologies.

Analyzing cellular populations in the early drug discovery process allows the complexity of living systems to be addressed and produces vast amounts of data that are more meaningful than those obtained from isolated proteins (Taylor, 2007). In combination with advanced bioinformatics tools treatments can be identified which lead to altered cell populations, and therefore might be relevant drugs or drug targets.

Nonetheless, several limitations in data analysis have restricted the full potential of high-throughput image-based assays so far (Lang, et al., 2006; Zhou and Wong, 2006). The usual course of events for a HCS analysis workflow starts with the extraction of image feature data, followed by normalization and statistical analysis, including final hit selection (Buchser, et al., 2004). A wide variety of microscopes, image-analysis and data-analysis software packages are available to address these issues (Gough and Johnston, 2007). However, distributions of multidimensional, multivariate phenotypic measurements from cellular populations are mostly transformed into single population-averaged values such as mean or median values. These population-averaged values are used for plate-wise or batch-wise normalizations, as well as for statistical analysis for hit selection (Birmingham, et al., 2009; Singh, et al., 2014), which leads to a substantial loss of information. Population-averaged values can indicate whether the value of the measured phenotype increases or decreases upon treatment, but do not reflect the detailed response of a cellular population to a certain treatment or gene depletion. Therefore, these population-averaged values are limiting the power of the statistical approaches that are widely used, such as Z-Score or percent-of-control (POC) analysis, making it impossible to identify more distinct reactions of a cell population. This loss of information also hampers the differentiation of treatments or gene depletions with the same ‘on-target’ effect from those with ‘off-target’ effects, which is extremely important for RNAi gene perturbation experiments, where multiple siRNAs are used per gene.

Some publications have described methods for non-averaged cell population data analysis from high-content image-based screens. Knapp et al. (Knapp, et al., 2011) showed considerable effects of population context on observed phenotypes when using non-averaged population data for the normalization steps, but still used population-averaged values for hit detection. Another method uses multivariate cell classification based on phenotypic changes for hit identification (Loo, et al., 2007), which results in a drug effect score, and a vector, indicating the simultaneous phenotypic changes induced by the drug. Another publication used multi-parametric phenotypic profiles to cluster genes based on morphological changes of individual cells (Fuchs, et al., 2010). Yet another group has proposed the use of Ripley’s K-function to identify knockdowns resulting in perturbation of this cell clustering (Suratanee, et al., 2010). Also the Kolomogorow-Smirnov (KS) test has been used to score the difference between control and samples populations (Gorenstein, et al., 2010). However, all these methods have limitations that prevent them from being widely used for large-scale high content cell population analysis. Multivariate classification methods are mostly based on the analysis of predominantly redundant image features, spatial clustering requires a subjective and work intensive classification step for the cellular populations and KS only uses one unique value to identify cell population with altered distributions.

Here we present a new approach called Single Object Profiles Regression Analysis (SOPRA) that overcomes many of these limitations by analyzing non-averaged cell population data. It uses a classification free regression analysis of normalized frequency distribution profiles of cell populations. SOPRA can be used to analyze data derived from various high-throughput techniques, such as images from automated microscopy or single cell data from FACS analysis. The regression workflow consists of i) a pre-processing step, ii) data gathering and normalization steps, iii) identification of significant profiles, iv) post-processing. The normalization is performed in a plate-wise and bin-wise fashion, resulting in a unique normalized frequency distribution profile for each feature of a cell population. Finally, normalized distribution profiles that exhibit statistically significant changes are identified by using a p-value and R-squared (RSQ)-value derived from the regression analysis with the R-package maSigPro (Conesa, et al., 2006). Additionally, normalized feature profiles that have been identified as significantly altered can be further clustered in a heatmap according to their similarity. This can be used to identify treatments with the same ‘on-target’ effects. Most loss-of-function screens use multiple siRNAs for the same gene, which should end up in the same cluster if they have a similar cell population phenotype. The more siRNAs for the same gene are identified as having a similar cell population profile, the more reliably this gene can be regarded as a hit. Beyond this, the derived values of a regression analysis of distribution profiles of cellular features are not affected by experimental bias to the same degree as population-averaged approaches (Sacher, et al., 2008), leading to more reproducible results. We used a cell-based chemical compound and RNAi screen of cell cycle progression to validate the SOPRA workflow. The cellular features ‘Area’, ‘Total Intensity DAPI’ and ‘Mean Intensity DAPI’ were extracted for each nucleus using image analysis software and subjected to the SOPRA workflow. We found that SOPRA can be used to identify statistically significant changes of frequency distribution profiles within cellular populations, whether induced by gene perturbation through siRNAs or by chemical inhibitor treatment. Taken together, SOPRA is a novel object-based data analysis workflow based on regression analysis of cellular feature distribution profiles to identify significantly changed cell populations from high-throughput data sets.

## Results

SOPRA utilizes a data gathering step combined with plate-wise and a so-called bin-wise normalization methods, as well as a two-step regression approach that first adjusts a global regression model with defined variables in order to identify profiles exhibiting statistically significant changes (Conesa, et al., 2006). The SOPRA workflow consists of several steps as outlined in Figure 1. *A: High Content Screen*. This first step includes screening, image analysis and data extraction. B: Preparation of screen description files and the single cell data files. *C: Preprocessing (optional)*. The derived data files for various image features at single cell level are subjected to a preprocessing step to exclude all data from flagged wells that should be excluded from the analysis. *D1: Data Gathering and Plate-Wise Normalization*. In this step each single cell object is annotated with additional information such as RNA.ID, plate number, well number, replicate number, well content and gene symbol. If the imaging software supports a gating procedure for objects that do not meet certain criteria, such as cell size, these can also be flagged and excluded from subsequent analysis steps. The measured value of each cell for the feature of interest is then normalized to the median of the objects in the neutral control wells. *D2: Data Gathering and Frequency Distribution Profiles (Histogram) Generation*. Next, the common binning axis of the distribution profiles is generated by determining the minimum and maximum limits of the measured feature across all the data of the screen to avoid strong relative differences at the tails of the distribution. The data are divided into equally spaced binning intervals, which is sufficient for population data that follows a given order of regression model (such as quadratic). A pseudo count of one is added to each bin to avoid bins with zero objects, and the relative frequency for each treatment is calculated by dividing the number of objects in each bin by the population size (sum of all objects in all bins). Next, a bin-wise normalization step is performed by dividing the relative frequency of each bin for each treatment by the median of the corresponding bin of the control wells, such as the ‘AllStars’ control.

**Figure 1.**
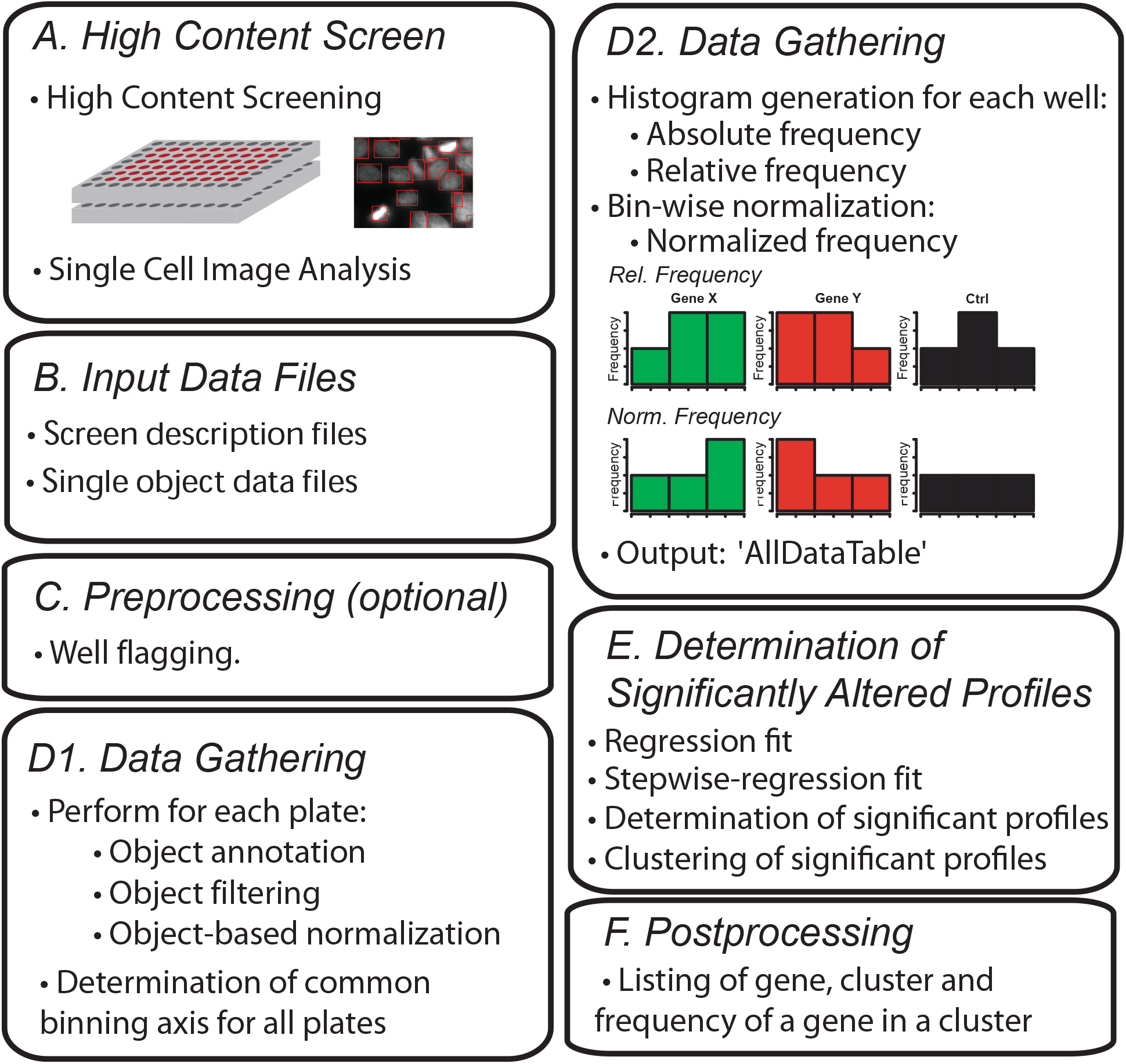
SOPRA workflow of high-throughput data sets. **(A)** High content screening data is generated and used to prepare single object data files and input data files. **(B)** Screen description and the single cell data files are generated manually. **(C)** Wells that should be omitted are flagged and **(D1)** the single object data is filtered, normalized to the median of the controls and a common binning axis for all plates is determined. **(D2)** For each measured feature the frequency distribution profile (histogram) is generated for each sample well, which is then normalized for each bin to the median distribution profile of the controls. **(E)** Significantly changed normalized distribution profiles are determined using regression analysis and **(F)** a post processing step is performed to determine the number of screening hits.

Binning creates equal-length bins to which data are assigned. The default number of bins (the binning level) is 7.

For variable x, assume that the data set is *{x*_*i*_*}*, let *x*_*1*_, *x*_*2*_,*…*.*x*_*m*_ represent the ordered values of the variable. Let the x^th^ percentile be *min(x)* and *max(x)*. The range of the variable is *range(x) = max(x) – min(x)*. For binning, the width of binning interval is 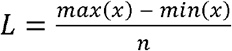. The split points are *s*_*k*_ *= min(x) + L * k*, where *k = 1, 2,…, numbin-1* and *numbin* is *n*. For each bin a pseudocount of 1 is added *Countp(X*_*ik*_ *) = Count(X*_*ik*_*) + 1*.

The output data, consisting of a normalized frequency distribution profile for each cell population and the annotation data are stored in the file ‘AllDataTable’. *E: Determination of Statistically Significant Altered Profiles*. A regression analysis is performed using the Bioconductor R-package maSigPro (Conesa, et al., 2006) to identify significantly changed normalized distribution profiles. *F: Postprocessing (optional)*. The gene, cluster and frequency of the gene within the cluster are listed for all significantly changed normalized distribution profiles identified by the maSigPro analysis.

To generate the data for the cell cycle progression screen we seeded HeLa cells in 384-well plates either transfected or treated with the inhibitors in three independent biological replicates (Figure 2A). We used 166 different siRNAs to target 107 genes, from which 54 had been reported to interfere with cell cycle progression (Kittler, et al., 2007) (Supplementary Figure 1, Supplementary Table 1). Additionally, we used cells treated with the chemical inhibitors aphidicolin or nocodazole, which lead to G1/S and G2/M cell cycle arrest, respectively. Cells left untreated (Mock) or treated with siRNAs against ‘AllStars’ or ‘Luciferase’ were used as negative controls. While images from AllStars- and Luciferase-treated cell populations showed an unaltered, normal phenotype, treatment of cells with aphidicolin (A1-4) or nocodazole (N1-4) resulted in an altered phenotype as a consequence of G1/S or G2/M phase arrest, respectively (Figure 2B). On day 4, cells were fixed, nuclei stained with Hoechst (Figure 2A), images acquired using automated microscopy and automated image analysis (Olympus Scan^R) was performed for extracting the image features ‘Area’, ‘Total Intensity DAPI’ and ‘Mean Intensity DAPI’ for each nucleus (Supplementary Figure 2) in tab-delimited files using a Scan^R export script. The cell population distribution profiles for the control as well as the chemically or siRNA-treated samples behave differently for the extracted object features (Figure 2C). They show a strong shift towards smaller nuclei for nocodazole, and towards larger nuclei for aphidicolin-treated samples for the feature ‘Area’, while for the feature ‘Mean Intensity DAPI’ the influence of these two chemical treatments on the mean intensity is the opposite. Interestingly, for the feature ‘Total Intensity DAPI’ a strong shift towards higher values was observed for nocodazole-treated samples, while aphidicolin treatment did not alter the profile compared to that of the ‘Allstars’, ‘Luciferase’ or ‘Mock’-treated wells. Distribution profiles of the cell populations treated with different siRNAs (samples) showed no clear tendency (Figure 2C).

**Figure 2.**
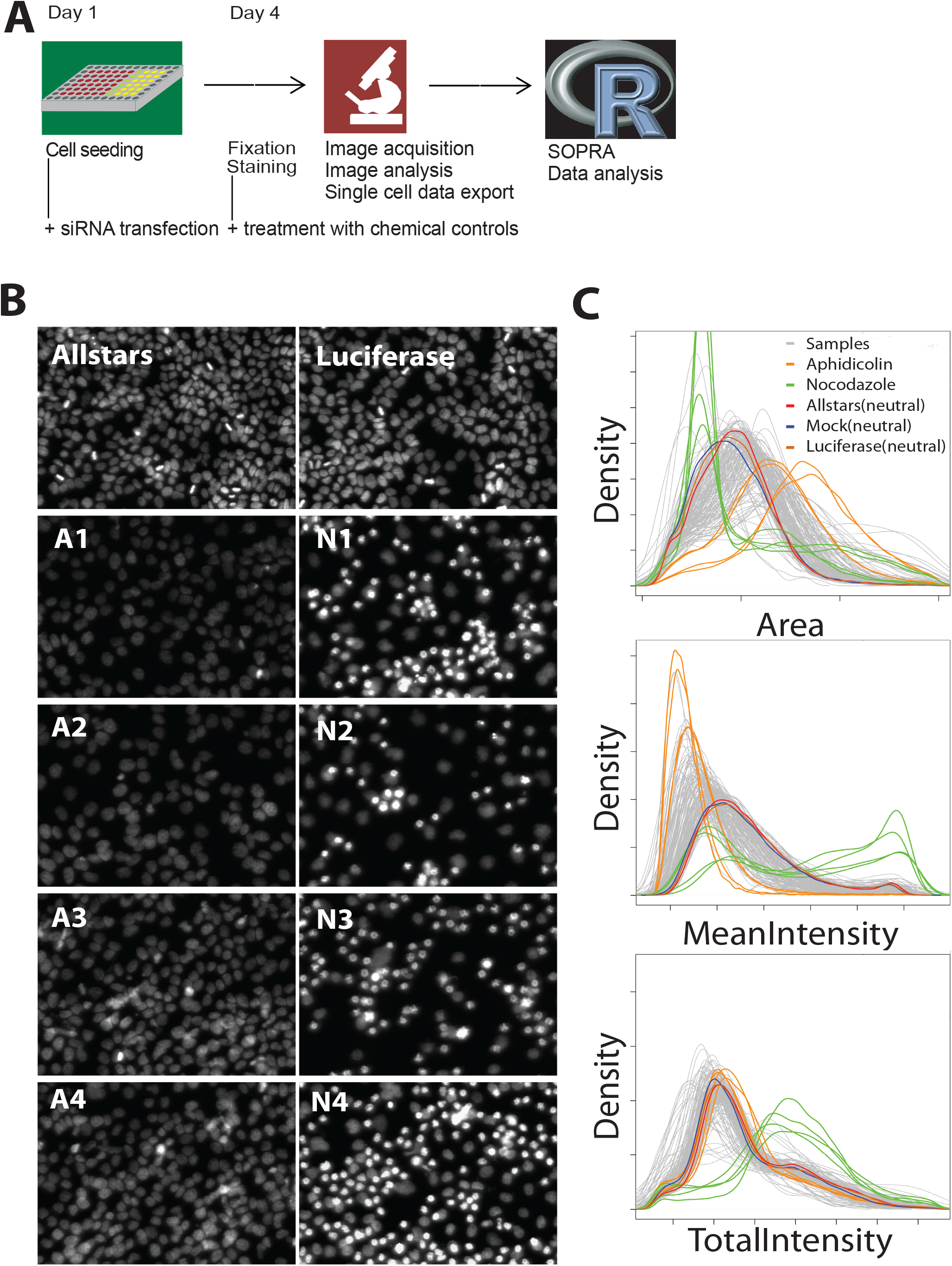
Schematic representation of the microscopic cell cycle screening assay. **(A)** Cells were seeded in 384-well plates and treated with siRNAs or chemical cell cycle inhibitors at different concentrations and time points to inhibit cell cycle progression. Cells were fixed, stained with Hoechst and subjected to automated microscopy and image analysis. **(B)** Treatment with the control-siRNAs AllStars and luciferase did not lead to any changes of the cell population. Treatment with aphidocoline A1 (2 μg/ml, 24 h), A2 (4 μg/ml, 24 h), A3 (2 μg/ml, 12 h), A4 (4 μg/ml, 12 h) and nocodazole N1 (50 ng/ml, 24 h), N2 (75 ng/ml, 24 h), N3 (50 ng/ml, 12 h), N4 (75 ng/ml, 12 h) resulted in cell populations arrested at various stages of the cell cycle. **(C)** Distribution profiles were generated for each well from the data exported for the features ‘Area’, ‘Mean Intensity DAPI’ and ‘Total Intensity DAPI’ for all nuclei.

We then calculated p-values and RSQ-values using maSigPro regression analysis, as described, to identify significantly altered distribution profiles compared to the neutral controls. The maSigPro package computes a regression fit for each frequency distribution profile, and uses a linear step-up (BH) false discovery rate (FDR) procedure (Benjamini and Hochberg, 1995). Here, we used a level of 0.05 for FDR control. Once statistically significant distribution profiles have been found, a variable selection procedure is applied to find significant variables for each profile. The final step is to generate lists of statistically significant profiles. As expected, cell populations treated with the ‘AllStars’ or ‘Luciferase’ controls usually had high p-values and low RSQ-values. Only two (10%) and four (20%) out of 20 cellular populations treated with the neutral controls ‘Allstars’ or ‘Luciferase’, respectively, were identified to be significantly changed for at least one of the three cellular features used (Supplementary Table 2; Figure 3A - Plate2, Well 207). When hits were only considered positive if at least two of the image features were identified as significantly changed, none of the neutral controls were identified as a hit. In contrast, cells treated with aphidicoline (A1-4) or nocodazole (N1-4) showed significant changes, indicated by low p-values and high RSQ-values for all of the three extracted cellular features (Figure 4A). All 28 profiles for each of the aphidicolin conditions A2 (4 μg/ml/24 h), A3 (2 μg/ml/12 h) and A4 (4 μg/ml/12 h), for each of the nocodazole conditions N1 (50 ng/ml/24 h), N2 (75 ng/ml/24 h), N3 (50 ng/ml/12 h) and N4 (75 ng/ml/12 h) and 27 out of 28 profiles for the aphidicolin condition A1 (2 μg/ml/24 h) were identified as significantly changed hits (Supplementary Figure 3). Interestingly, aphidicolin-treated samples showed marked differences for the cellular features ‘Area’ and ‘Mean Intensity’ and only slight changes for the cell feature ‘Total Intensity DAPI’ (Figure 3A – A1: Plate1, Well 208, A2: Plate2, Well 353, A3: Plate1, Well 44, A4: Plate2, Well 213), while nocodazole-treated samples showed strong changes in all three cellular features used (Figure 3A -N4: Plate1, Well 47, N1: Plate 2, Well 354). In total, using these thresholds for the p-value and the RSQ-value, 359 normalized distribution profiles were identified as significantly altered for each of the cellular features ‘Area’ and ‘Mean Intensity DAPI’ and 335 normalized distribution profiles for the cellular feature ‘Total Intensity DAPI’. This resulted in a total of 448 significantly changed cell populations; with 247 profiles significantly changed for all three, 111 profiles for two and 90 profiles for only one of the analyzed cellular features. Next, for the 448 profiles identified as significantly changed, a k-means clustering approach was performed (Figure 4B and 4C). The normalized distribution profiles for the features ‘Area’, ‘Mean Intensity DAPI’ and ‘Total Intensity DAPI’ were arranged in four, three and two profile clusters, respectively. Cluster numbers were selected to give high cluster reproducibility. Finally, for all clustered profiles a heatmap was defined, based on the k-means clustering result arranged as a vector (consisting of zeros and ones such as 0001-010-10 for a profile resulting in cluster 4 for ‘Area’, cluster 2 for ‘Mean Intensity DAPI’ and cluster 1 for ‘Total Intensity DAPI’). The heatmap was sorted using a hierarchical clustering (hclust) algorithm to identify cell populations with similar distribution profiles (Figure 3A). Finally, a dendrogram cut-off value of 1.8 was used to generate three main groups in the matrix.

**Figure 3.**
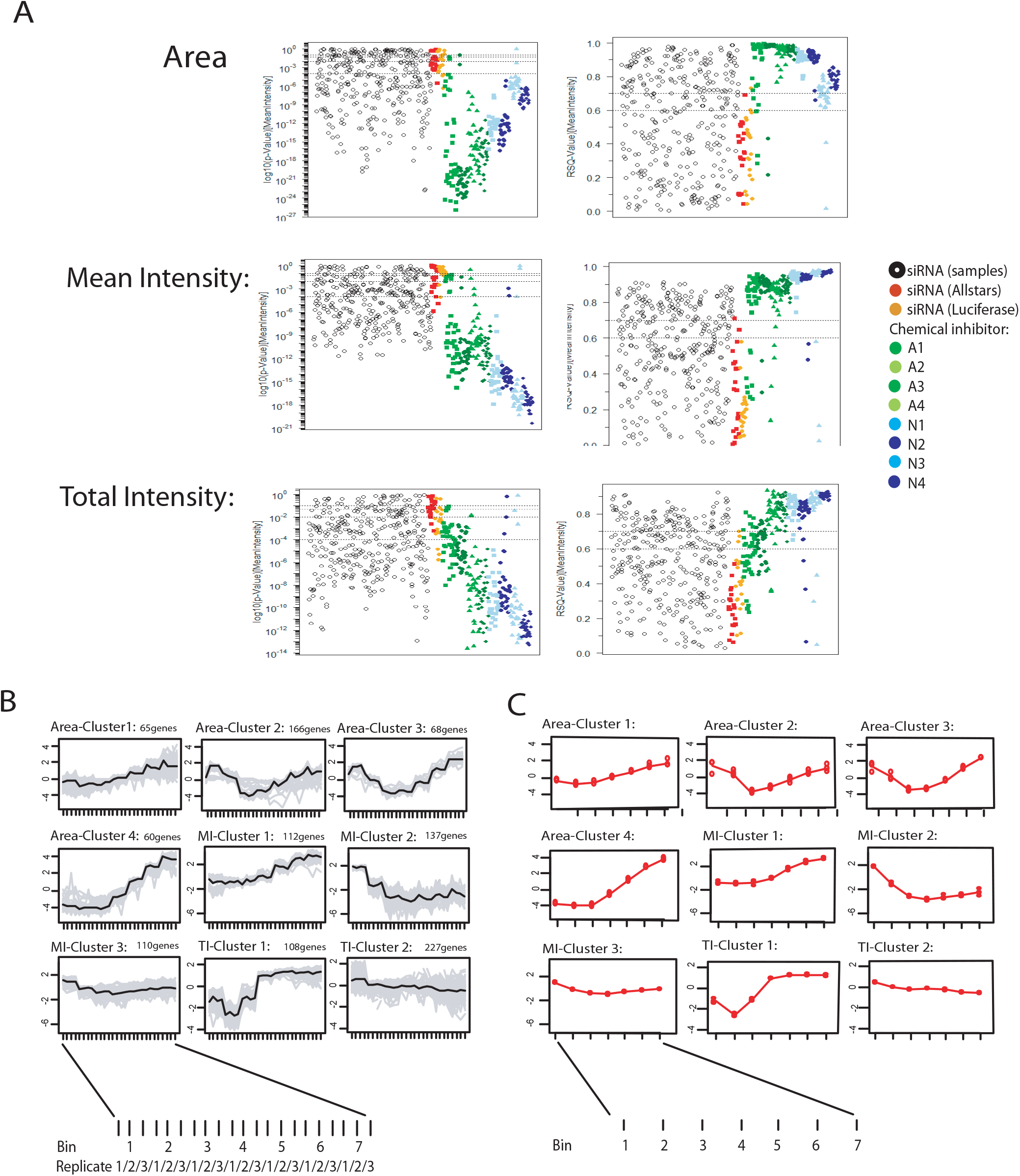
Heatmap analysis and examples of significantly altered distribution profiles. **(A)** The normalized regression profiles for different treatment conditions for aphidicolin (A1-A4) and nocodazole (N1 and N4), as well as Luciferase are displayed. A heatmap was generated showing the distribution of all cell populations with at least one significantly changed profile for the features ‘Area’, ‘Mean Intensity DAPI’ and ‘Total Intensity DAPI’ among the SOPRA cluster profiles. Wells treated with aphidicolin or nocodazole are displayed in different shades of green or blue in the row sidebar. Wells Mock-treated or treated with siRNA against Luciferase or AllStars are indicated in red, orange and yellow, respectively. Wells treated with siRNA against specific genes are displayed in grey in the row sidebar. The heatmap is clustered using hierarchical clustering, and a dendogram, cut-off of 1.8 performed resulting in the heatmap groups (1), (2) and (3). The Venn diagram displays the distribution of the significantly changed profiles for each treatment among the heatmap groups (1)-(3). **(B)** Examples of profiles for the features ‘Area’, ‘Total Intensity DAPI’ and ‘Mean Intensity DAPI’ of cell populations significantly changed upon siRNA treatment, as well as the corresponding microscopic and FACS images.

**Figure 4.**
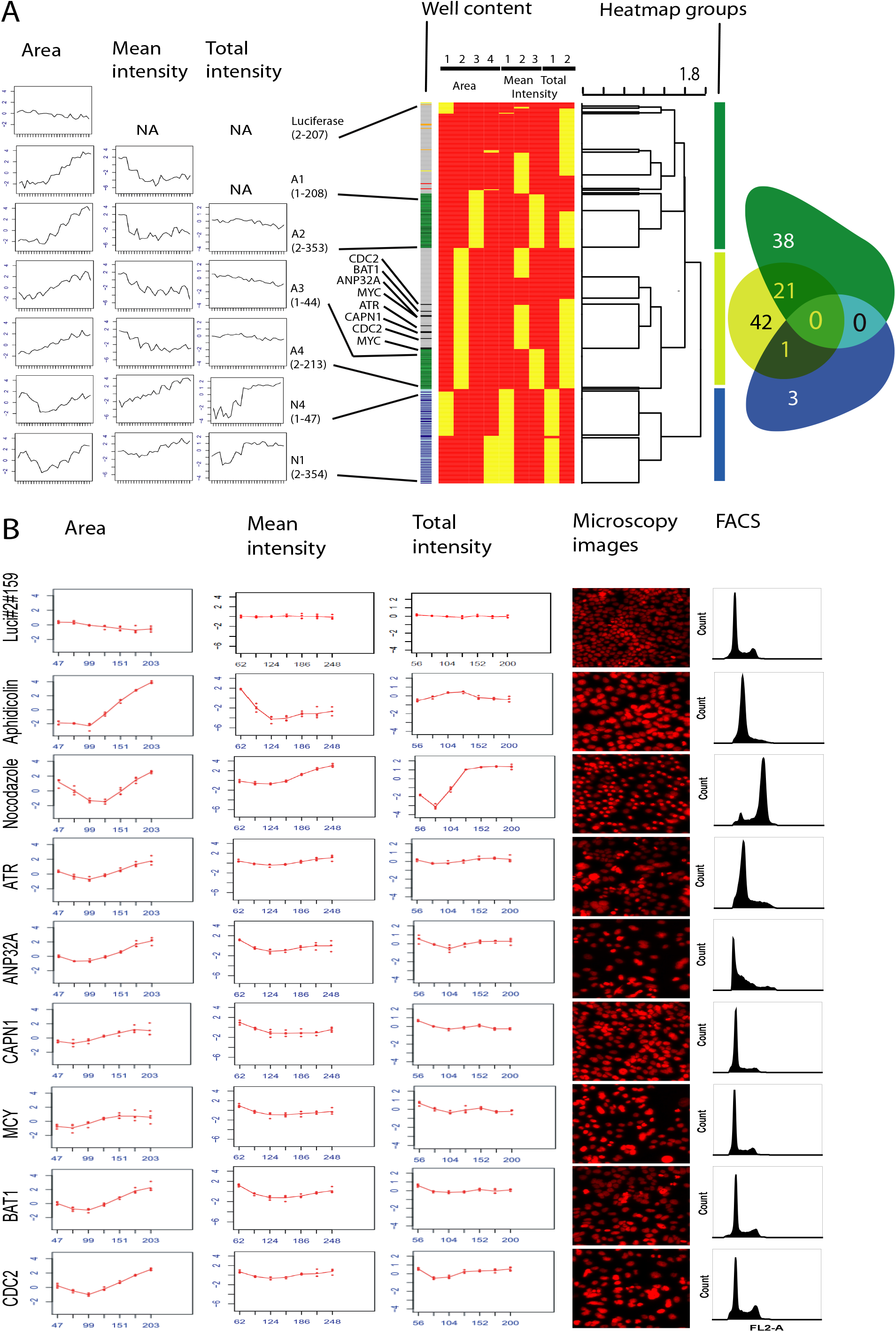
Results of SOPRA regression analysis and cluster profiles. **(A)** Calculated RSQ and p-values of each well for the features ‘Area’, ‘Mean Intensity DAPI’ and ‘Total Intensity DAPI’ using the maSigPro package. **(B)** Data visualization by cluster analysis. Normalized distribution profiles of all significantly altered normalized profiles for the three image features were clustered using k-means with 4, 3 and 2 clusters, respectively. The average feature profile is shown (black line) together with the individual profiles of the cell populations in the cluster (grey lines) or **(C)** as the mean of 3 replicates.

As a result, the aphidicolin-treated samples A1 and A2 grouped differently from the aphidicolin-treated samples A3 and A4, (Figure 3A, sidebar), while the nocodazole-treated samples N1 and N2, as well as N3 and N4, grouped together. Further, the significant distribution profiles of samples treated with siRNA were more dispersed in the heatmap, depending on the individual feature distribution. Thus, with this two-step method - first identifying statistically significant normalized profiles for each analyzed image feature, then using a heatmap to generate profile groups – we were able to differentiate cell populations showing a similar distribution among the cluster profiles. Taken together, the SOPRA workflow was responsive enough to distinguish not only nocodazole-treated from aphidicolin-treated samples, but also to differentiate between samples that were treated with the same concentration but for different durations (A1/A2 vs A3/A4).

As laid out above, analyzed features for siRNAs that target the same gene and have the same ‘on-target’ phenotype should end up in the same cluster and also in the same heatmap group. Therefore, we further analyzed if individual siRNAs for the same gene were represented in the same or different heatmap groups. The individual siRNAs of 38 and 42 genes appeared exclusively in profile groups 1 or 2, respectively, strengthening the ‘on-target’ specificity of these siRNAs. In contrast, individual siRNAs of 21 genes were represented in both of these groups, indicating less stringent ‘on-target’ specificity or other influences, such as experimental variation. For the SOPRA workflow, two different siRNAs were used for each gene in duplicate, therefore hits were classified as medium or weak hits if the two siRNAs did not show the same cluster profile and were not grouped in the same heatmap group.

To assess the reproducibility of plate replicates and SOPRA workflow the RSQ-values for different groups of replicates were determined and the correlation matrix between these groups was calculated. Firstly, we defined replicate group R1 containing replicate r2, r3 and r4 (i.e. without replicate 1), replicate group R2 containing r1, r3, and r4 (without replicate 2) and so on. Running SOPRA for cell feature ‘Total Intensity DAPI’ with pre-defined maSigPro parameters alfa=1, Q=1 and RSQ=0 one gets the p-values and RSQ-values for each replicate group. The correlation matrix between the RSQ-values for sample data from the different groups R1-R4 is calculated and visualized (Figure 5).

**Figure 5.**
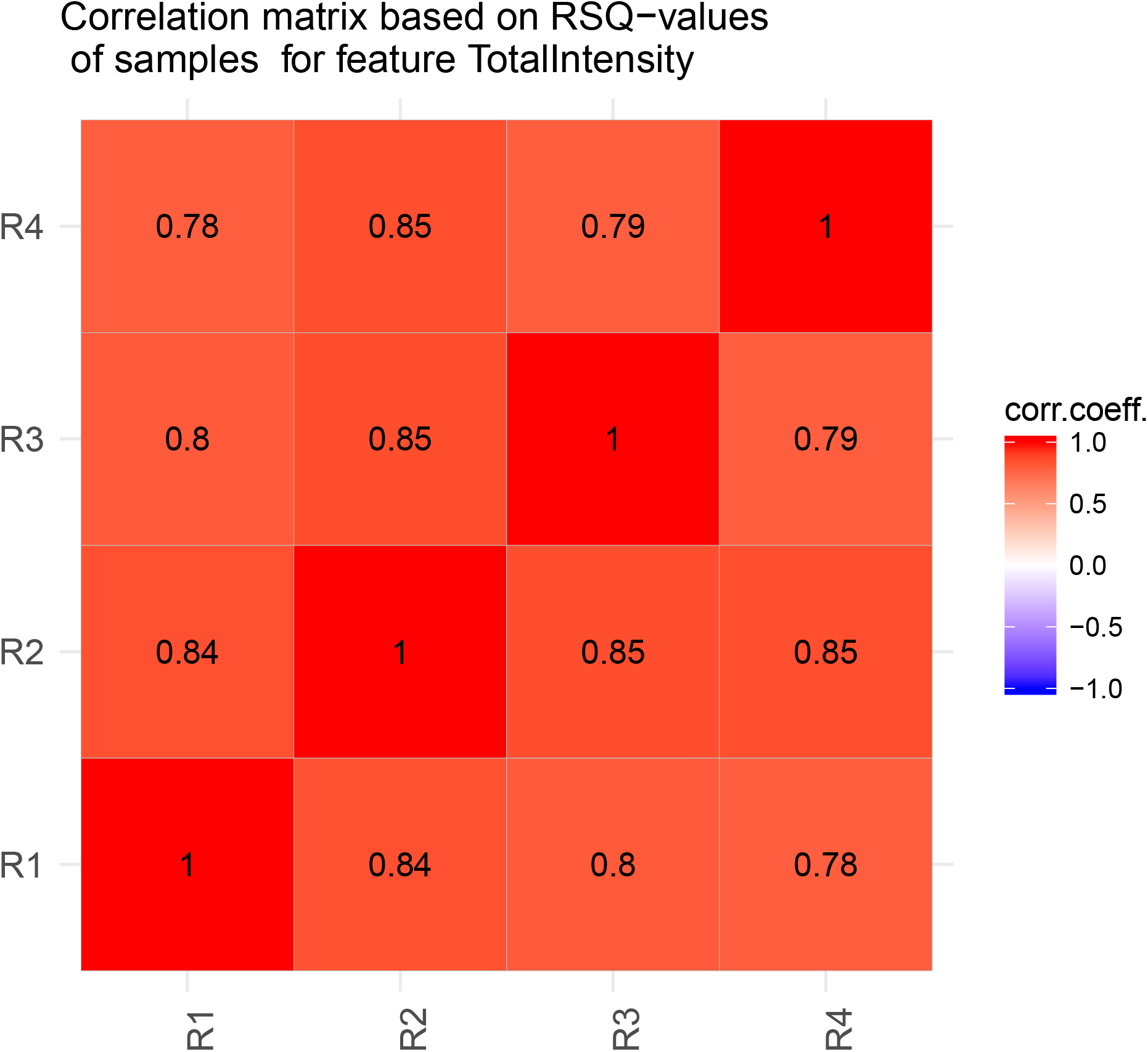
Reproducibility assessment between replicates. Correlation matrix between the RSQ-values for sample data from the different replicate groups R1-R4 for cell feature ‘Total Intensity DAPI’

Furthermore, we used Receiver Operating Characteristic (ROC) to assess the statistical performance of SOPRA workflow in comparison to other approach, such as “Kolmogorov-Smirnov (KS) test” which uses probability density and “t.test” for assessment of population differences. The RO curve for cell feature ‘Total Intensity DAPI’ is depicted in Figure 6 and show that the SOPRA method lies between the other two RO curves.

**Figure 6.**
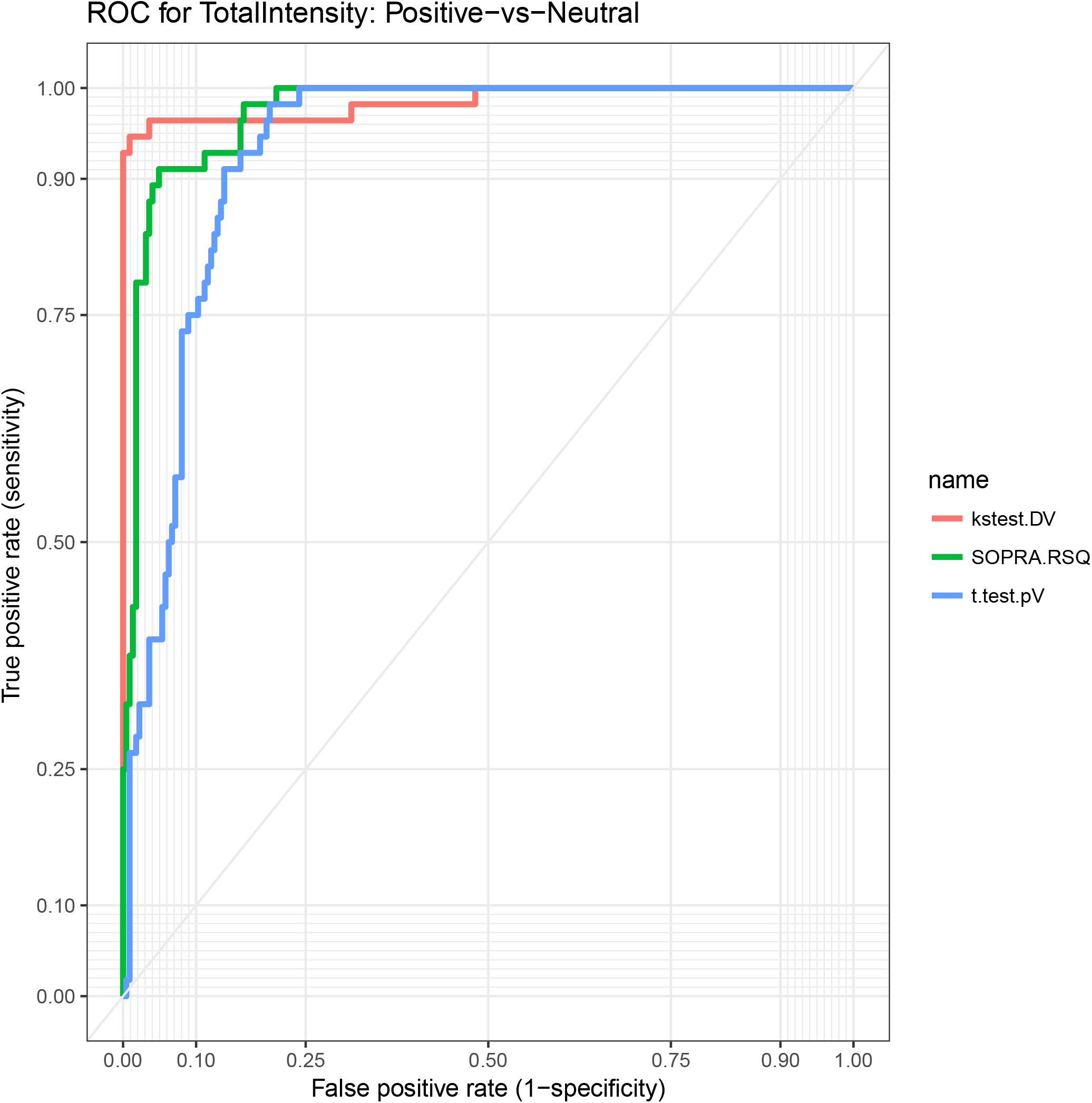
Assessment of diagnostic quality by Reciever Operating Curve (ROC) Receiver Operating Characteristic (ROC) serves to assess the SOPRA workflow in comparison to other statistical approaches, such as “t.test” and “Kolmogorov-Smirnov (KS) test”. The RO curves for cell feature ‘Total Intensity DAPI’ shows that the SOPRA method lies between the t.test and KS-test.

To benchmark the efficiency of this method in gene perturbation hit prediction, we tested whether the results of the SOPRA workflow could be validated by either the original cell cycle data from Kittler *et al*. (Kittler, et al., 2007) or FACS data generated by our group (Figure 3B). We selected 46 of the genes (hits and non-hits) analyzed with SOPRA and performed FACS analysis for cell cycle profiles with one siRNA per gene. A hit was scored as positive for a particular method if at least one other method also leads to the same (positive or negative) result (Supplementary Table 3). Out of the 46 genes analyzed, 30 genes from the Kittler *et al*. study were validated with at least one of the other methods (SOPRA or FACS), while for the SOPRA and FACS analyses 36 and 38 genes, respectively, were validated by one of the other two methods. Taken together, SOPRA and FACS analysis scored best in their ability to predict hits, compared to the data published by Kittler *et al*. (Kittler, et al., 2007).

Thus, the SOPRA workflow offers a unique and fast analysis approach, based on measured single features of cell populations, comparable to or better than published methods. In contrast to FACS data analysis it does not need manual intervention or thresholding, such as cell gating. SOPRA is therefore well suited for high-throughput and high-content data, as it can be easily run on multiple features from an identical cell population.

## Conclusions

Most methods published for analyzing high-content microscopic screens use population-averaged values or manually performed cell classification steps for normalization and hit classification. The SOPRA workflow represents a novel approach for analyzing large microscopy-based high-content screens using non-averaged data of cell populations for normalization and hit determination. The workflow generates frequency distribution profiles of cellular features normalized to a neutral control for each treatment. These normalized distribution profiles are used for hit identification by regression analysis to identify significantly altered profiles using the R-package maSigPro, as originally described for the analysis of single series time course gene-expression data.

RNAi screens are frequently performed with multiple siRNAs per target gene; however the use of population-averaged values often leads to the identification of ‘off-target’ effects as hits, since population averaged values can only monitor major variations of the phenotype such as up- or down-regulation compared to a control. In contrast, non-averaged data can indicate more diverse changes of a cell population upon treatment; thus different siRNAs targeting the same gene should have a similar ‘on-target’ effect on the distribution profile of the measured cellular features and consequently these are more likely to be ‘true’ hits. The SOPRA workflow we describe here has the power to cluster all significantly altered normalized distribution profiles, identifying siRNAs with similar ‘on-target’ profiles for the same gene via a heatmap approach. Therefore, the SOPRA workflow can be used to avoid false-positive hits or ‘off-target’ effects, leading to more reliable HCS hit results, reducing time and work intensive validation steps.

In principle, the SOPRA workflow can be used to analyze single cell population data from various sources such as microscopy or FACS. In this study, we performed a microscopy-based high content screen of the effect of siRNA-mediated gene knockdown of selected genes taken from a published cell cycle data set from Kittler et al (Kittler, et al., 2007), as an example to demonstrate the utility of the SOPRA workflow.

We were able to show that the false positive detection rate (detection of neutral controls as significantly changed) can be reduced considerably when taking into account more than one cellular feature. As described using the generated cell cycle data, we were able to demonstrate that the SOPRA workflow led to no false-positive hits among the neutral controls, when at least two of the image features were taken into account. For the cell populations treated with the cell cycle inhibitors, a very high hit detection rate of 99.55% was achieved (223 of 224 cell population profiles). We also used siRNA knockdowns in this screen, which produce less significant phenotypic effects compared to small chemical compounds. Nevertheless, analysis of changed cell populations based on gene perturbation with siRNA using SOPRA still achieved a hit detection rate comparable to a manual FACS analysis with commercial software, which requires predetermined gating or thresholding.

Taken together, SOPRA is a novel analysis workflow that uses a unique analysis approach for non-averaged high-throughput data from cellular features, based on regression analysis of normalized frequency distribution profiles of cell populations. It offers an easy to handle workflow and can be run on hundreds of cell populations using multiple features. In particular, treated cell populations are defined as significantly changed on two measurements - the p-value and the RSQ-value - followed by a clustering step to identify treatments with the same normalized density profiles. A following heatmap analysis enabled us to filter out most hits that are likely to be false positive. Thus, SOPRA is a unique tool ideal for high content analysis of cell population data.

## Methods

### Cell Cycle perturbation screen

We generated a set of screening plates consisting of siRNAs (Qiagen, Germany) targeting proteins responsible and not responsible for cell cycle progression, as well as the neutral siRNAs ‘AllStars’ and ‘Luciferase’, and wells without treatment (Mock) (Supplementary Figure 1). On day one cells were seeded in 96-well plates and transfected using Hiperfect (Qiagen, Germany). The chemical cell cycle inhibitors nocodazole and aphidicolin were added as positive controls at the described time points and concentrations. On day four cells were fixed using 4% PFA and stained with Hoechst 33342 (5 μg/ml, Sigma). The plates were imaged using an automated microscope (IX-81, Olympus, Germany) and analyzed using the Scan^R software with an image analysis assay designed in-house (Supplementary Figure 2).

Using a Scan^R single cell export script, single cell data was exported and are downloadable from https://transfer.mpiib-berlin.mpg.de/s/AibR4AHLCR9xzDB?path=%2F. The SOPRA project description (Supplementary File 1) is also available from GitHub https://github.com/kppleissner/SOPRA/.

### Cell Cycle FACS validation

For FACS analysis of cell cycle profiles, 1 × 10^5^ cells were seeded into each well of a 12-well plate 24 h before transfection. Cells were then transfected with Hiperfect transfection reagent (Qiagen) according to the manufacturer’s guidelines. In brief, 150 ng of specific siRNA was added to RPMI without serum and incubated with 6 μl Hiperfect in a total volume of 100 μl. After 10 to 15 min, the liposome-siRNA mixture was added to the cells with 1 ml of cell culture medium (RPMI (Gibco) supplemented with 10% fetal calf serum (FCS) (Biochrome), 2 mM glutamine, and 1 mM sodium pyruvate), to give a final siRNA concentration of 10 nM. After 1 day, cells were trypsinized and seeded into new 6-well plates. Three days after transfection, cells were detached from the plate with the addition of trypsin-EDTA for 5 minutes, spun down for 10 minutes at 500 x *g* and resuspended in 0.5 ml PBS. The resuspended cells were then added to 70% ethanol for fixation and left at –20°C overnight. Cells were collected by centrifugation, resuspended, rinsed in PBS and re-collected by centrifugation. Pelleted cells were resuspended in 500 μl PBS containing a final concentration of 20 μg/ml propidium iodide and 200 μg/ml RNAse A and left in the dark for 30 minutes at room temperature. Cell Cycle analysis was then performed using a Becton Dickinson FACsort flow cytometer and BD CellQuest Pro Software (BD Biosciences).

### SOPRA

The SOPRA workflow (Figure 1) consists of several steps and requires a variety of input files. The *‘Single Cell Feature Files’* contain the features for every single cell measured, while the files *‘PlateConf_LookUp’, ‘PlateList’* and *‘ScreenLog’* contain information about well content, plate content and flagged wells. In the first step, the data is gathered, including flagging of wells and single objects within wells. In the next step, a plate-wise median normalization is performed and the limits for the binning intervals are defined. Subsequently, the single objects within each binning interval (bin) are counted, and a bin-wise normalization is performed. Derived frequency distribution profiles of measured features are then subjected to the regression analysis using R-Package maSigPro. Significantly different profiles can be identified using the calculated p-value, RSQ-value and alpha-value for each sample profile. The significant profiles can be clustered using different clustering algorithms. Finally, a post processing step (optional) can be performed in order to convert siRNA into gene names, cluster membership and frequency. The SOPRA workflow is written as a <u>Shiny application in R</u>. A detailed project description with specific instructions for how to run the workflow is available from GitHub.

## Code availability and implementation

Source code of SOPRA shiny application (ui.R, server.R), single cell data (96-wells plate) data for testing and SOPRA project description (folder: Manual) are freely available from GitHub https://github.com/kppleissner/SOPRA/.

## Supporting Data ZIP File for 384-wells plates (Single Cell Features)

Due to large size of files the 384-wells plate data couldn’t be uploaded to GitHub and therefore are available as ***384_Plates_for_SOPRA*.*zip*** from MPI-IB Cloud tranfer server via this URL https://transfer.mpiib-berlin.mpg.de/s/AibR4AHLCR9xzDB?path=%2F.

The ***384_wells_Plates_for_SOPRA*.*zip*** file contains data based on a cell cycle screen analyzed with Scan^R (Olympus). Following cell features were measured: ‘Area’, ‘Mean Instensity DAPI’ and ‘Total Intensity DAPI’. In general, any file of the correct format can be used for SOPRA. For each plate – or part of a plate – one file is needed. The folders also contain the descriptive files *‘PlateConf_LookUp’, ‘PlateList’* and *‘ScreenLog’*.

## Author contributions

RKG-Conceived, designed, performed and analyzed the screen and wrote the manuscript

KPP-Wrote R-Scripts for SOPRA workflow, replication and ROC analysis, shiny interface, project description and realized storage of SOPRA into GitHub

CC – Performed the FACS validation of hits

TFM – Supervised the project

APM-Conceived the project, conceived, designed and analyzed the screen, wrote the R-Scripts, user interface UI.R and server.R in the Shiny environment, the SOPRA-Project Description and the manuscript.

## Acknowledgements

The authors would like to thank Kathrin Lättig for excellent technical support, Oliver Friedrichs and Ralf Träger for IT support, Hilmar Berger for critical reading and Rike Zietlow for editing the manuscript.

This work was supported by the Bundesministerium für Wirtschaft und Energie (Federal Ministry for Economic Affairs and Energy) BMWi ZIM program (grant no. KF3149632LW4).

