## Supplementary Figure 1 for "Single Object Profiles Regression Analysis (SOPRA): A novel method for analyzing high content cell-based screens": Supplementary Figure 1.pdf

Plate1

| | 1 | 2 | 3 | 4 | 5 | 6 | 7 | 8 | 9 | 10 | 11 | 12 | 13 | 14 | 15 | 16 | 17 | 18 | 19 | 20 | 21 | 22 | 23 | 24 |
|---|---|---|---|---|---|---|---|---|---|---|---|---|---|---|---|---|---|---|---|---|---|---|---|---|
| A | | | | | | | | | | | | | | | | | | | | | | | | |
| B | | 1 | 1 | 15 | 15 | 29 | 29 | 43 | 43 | 57 | 57 | 71 | 71 | AS | M | A1 | A2 | N1 | N2 | A3 | A4 | N3 | N4 | Plk 1-1μM |
| C | | 2 | 2 | 16 | 16 | 30 | 30 | 44 | 44 | 58 | 58 | 72 | 72 | AS | M | A1 | A2 | N1 | N2 | A3 | A4 | N3 | N4 | Plk 1-1μM |
| D | | 3 | 3 | 17 | 17 | 31 | 31 | 45 | 45 | 59 | 59 | 73 | 73 | AS | M | A1 | A2 | N1 | N2 | A3 | A4 | N3 | N4 | Plk 1-200nM |
| E | | 4 | 4 | 18 | 18 | 32 | 32 | 46 | 46 | 60 | 60 | 74 | 74 | AS | M | A1 | A2 | N1 | N2 | A3 | A4 | N3 | N4 | Plk 1-200nM |
| F | | 5 | 5 | 19 | 19 | 33 | 33 | 47 | 47 | 61 | 61 | 75 | 75 | AS | L | A1 | A2 | N1 | N2 | A3 | A4 | N3 | N4 | Plk 1-40nM |
| G | | 6 | 6 | 20 | 20 | 34 | 34 | 48 | 48 | 62 | 62 | 76 | 76 | L | L | A1 | A2 | N1 | N2 | A3 | A4 | N3 | N4 | Plk 1-40nM |
| H | | 7 | 7 | 21 | 21 | 35 | 35 | 49 | 49 | 63 | 63 | 77 | 77 | L | L | A1 | A2 | N1 | N2 | A3 | A4 | N3 | N4 | Plk 1-8nM |
| I | | 8 | 8 | 22 | 22 | 36 | 36 | 50 | 50 | 64 | 64 | 78 | 78 | L | L | A1 | A2 | N1 | N2 | A3 | A4 | N3 | N4 | Plk 1-8nM |
| J | | 9 | 9 | 23 | 23 | 37 | 37 | 51 | 51 | 65 | 65 | 79 | 79 | L | L | A1 | A2 | N1 | N2 | A3 | A4 | N3 | N4 | Plk 1-1.6nM |
| K | | 10 | 10 | 24 | 24 | 38 | 38 | | | 66 | 66 | 80 | 80 | L | AS | A1 | A2 | N1 | N2 | A3 | A4 | N3 | N4 | Plk 1-1.6nM |
| L | | 11 | 11 | 25 | 25 | 39 | 39 | 53 | 53 | 67 | 67 | 81 | 81 | M | AS | A1 | A2 | N1 | N2 | A3 | A4 | N3 | N4 | |
| M | | 12 | 12 | 26 | 26 | 40 | 40 | 54 | 54 | 68 | 68 | 82 | 82 | M | AS | A1 | A2 | N1 | N2 | A3 | A4 | N3 | N4 | |
| N | | 13 | 13 | 27 | 27 | 41 | 41 | 55 | 55 | 69 | 69 | 83 | 83 | M | AS | A1 | A2 | N1 | N2 | A3 | A4 | N3 | N4 | |
| O | | 14 | 14 | 28 | 28 | 42 | 42 | 56 | 56 | 70 | 70 | 84 | 84 | M | AS | A1 | A2 | N1 | N2 | A3 | A4 | N3 | N4 | |
| P | | | | | | | | | | | | | | | | | | | | | | | | |

Legend for controls

| |
|---|
| AS-AllStars |
| L-Luci |
| M-mock |
| A1- Aphidicolin- 2ug/ml- 24hr |
| A2- Aphidicolin- 4ug/ml- 24hr |
| A3- Aphidicolin- 2ug/ml- 12hr |
| A4-Aphidicolin- 4ug/ml- 12hr |
| N1-Nocodazole- 50ng/ml- 24hr |
| N2-Nocodazole- 75ng/ml- 24hr |
| N3-Nocodazole- 50ng/ml- 12hr |
| N4-Nocodazole- 75ng/ml- 12hr |
| Test siRNA's |
| PLK1-transfection control |
| empty |

Plate2

| | 1 | 2 | 3 | 4 | 5 | 6 | 7 | 8 | 9 | 10 | 11 | 12 | 13 | 14 | 15 | 16 | 17 | 18 | 19 | 20 | 21 | 22 | 23 | 24 |
|---|---|---|---|---|---|---|---|---|---|---|---|---|---|---|---|---|---|---|---|---|---|---|---|---|
| A | | | | | | | | | | | | | | | | | | | | | | | | |
| B | | 85 | 85 | 99 | 99 | 113 | 113 | 127 | 127 | 141 | 141 | 155 | 155 | AS | M | A1 | A2 | N1 | N2 | A3 | A4 | N3 | N4 | Plk 1-1μM |
| C | | 86 | 86 | 100 | 100 | 114 | 114 | 128 | 128 | 142 | 142 | 156 | 156 | AS | M | A1 | A2 | N1 | N2 | A3 | A4 | N3 | N4 | Plk 1-1μM |
| D | | 87 | 87 | 101 | 101 | 115 | 115 | 129 | 129 | 143 | 143 | 157 | 157 | AS | M | A1 | A2 | N1 | N2 | A3 | A4 | N3 | N4 | Plk 1-200nM |
| E | | 88 | 88 | 102 | 102 | 116 | 116 | 130 | 130 | 144 | 144 | 158 | 158 | AS | M | A1 | A2 | N1 | N2 | A3 | A4 | N3 | N4 | Plk 1-200nM |
| F | | 89 | 89 | 103 | 103 | 117 | 117 | 131 | 131 | 145 | 145 | 159 | 159 | AS | L | A1 | A2 | N1 | N2 | A3 | A4 | N3 | N4 | Plk 1-40nM |
| G | | 90 | 90 | 104 | 104 | 118 | 118 | 132 | 132 | 146 | 146 | 160 | 160 | L | L | A1 | A2 | N1 | N2 | A3 | A4 | N3 | N4 | Plk 1-40nM |
| H | | 91 | 91 | 105 | 105 | 119 | 119 | 133 | 133 | 147 | 147 | 161 | 161 | L | L | A1 | A2 | N1 | N2 | A3 | A4 | N3 | N4 | Plk 1-8nM |
| I | | 92 | 92 | 106 | 106 | 120 | 120 | 134 | 134 | 148 | 148 | 162 | 162 | L | L | A1 | A2 | N1 | N2 | A3 | A4 | N3 | N4 | Plk 1-8nM |
| J | | 93 | 93 | 107 | 107 | 121 | 121 | 135 | 135 | 149 | 149 | 163 | 163 | L | L | A1 | A2 | N1 | N2 | A3 | A4 | N3 | N4 | Plk 1-1.6nM |
| K | | 94 | 94 | 108 | 108 | 122 | 122 | 136 | 136 | 150 | 150 | 164 | 164 | L | AS | A1 | A2 | N1 | N2 | A3 | A4 | N3 | N4 | Plk 1-1.6nM |
| L | | 95 | 95 | 109 | 109 | 123 | 123 | 137 | 137 | 151 | 151 | 165 | 165 | M | AS | A1 | A2 | N1 | N2 | A3 | A4 | N3 | N4 | |
| M | | 96 | 96 | 110 | 110 | 124 | 124 | 138 | 138 | 152 | 152 | 166 | 166 | M | AS | A1 | A2 | N1 | N2 | A3 | A4 | N3 | N4 | |
| N | | 97 | 97 | 111 | 111 | 125 | 125 | 139 | 139 | 153 | 153 | 167 | 167 | M | AS | A1 | A2 | N1 | N2 | A3 | A4 | N3 | N4 | |
| O | | 98 | 98 | 112 | 112 | 126 | 126 | 140 | 140 | 154 | 154 | | | M | AS | A1 | A2 | N1 | N2 | A3 | A4 | N3 | N4 | |
| P | | | | | | | | | | | | | | | | | | | | | | | | |
