## Supplementary figures and images for "Single Object Profiles Regression Analysis (SOPRA): A novel method for analyzing high content cell-based screens"

### Supplementary Figure 2.pdf

#### Supplementary Figure 2

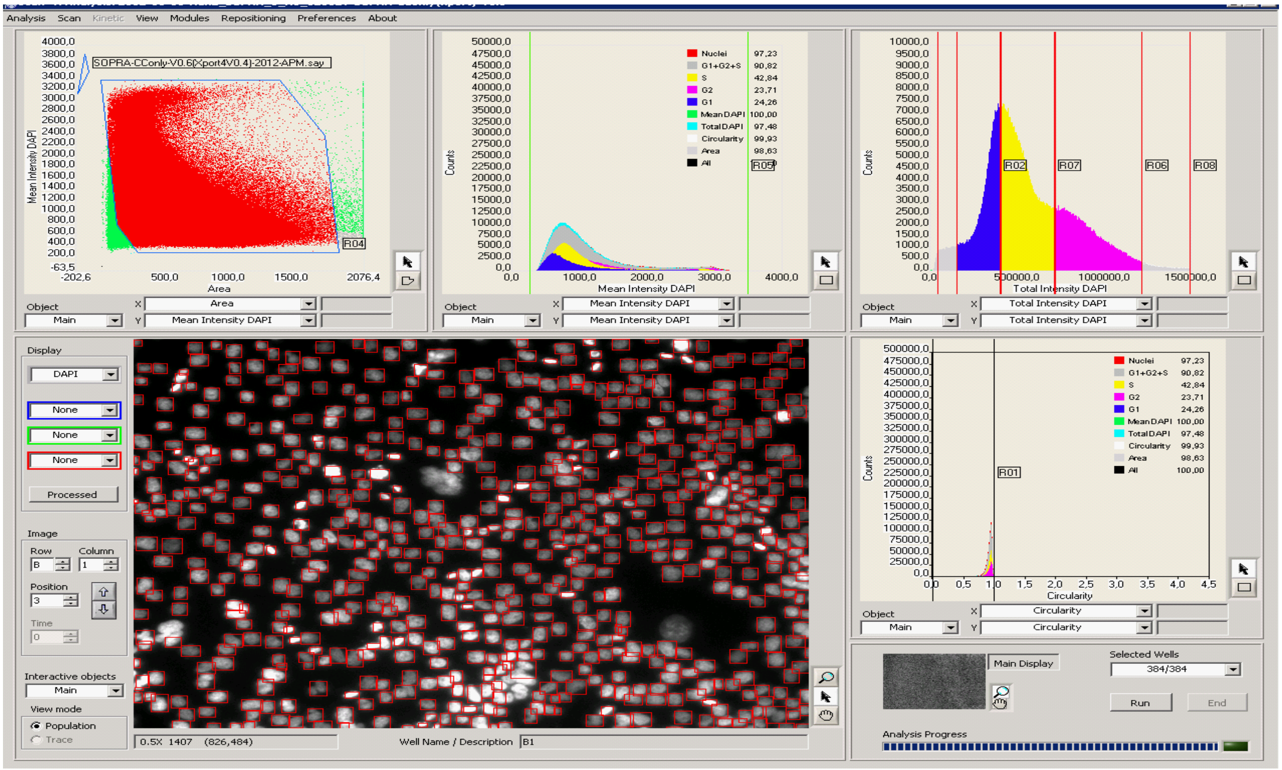
