## Supplementary Figure 3 for "Single Object Profiles Regression Analysis (SOPRA): A novel method for analyzing high content cell-based screens": Supplementary Figure 3.pdf

| Treatment | | | % |
|---|---|---|---|
| ***A1*** | 27 | 28 | 96.43 |
| ***A2*** | 28 | 28 | 100.00 |
| ***A3*** | 28 | 28 | 100.00 |
| ***A4*** | 28 | 28 | 100.00 |
| ***N1*** | 28 | 28 | 100.00 |
| ***N2*** | 28 | 28 | 100.00 |
| ***N3*** | 28 | 28 | 100.00 |
| ***N4*** | 28 | 28 | 100.00 |
| ***Luci*** | 4 | 20 | 20.00 |
| ***Mock*** | 3 | 18 | 16.67 |
| ***Allstars*** | 2 | 20 | 10.00 |

| | *Number of ..* | | *Per cent of ..* |
|---|---|---|---|
| | *..genes as hits or nonhits* | *..genes validated by at least one other method* | |
| SOPRA hits | 35 | 29 | 83% |
| SOPRA nonhits | 9 | 7 | 78% |
| FACS hits | 33 | 29 | 88% |
| FACS nonhits | 11 | 9 | 82% |
| Paper hits | 19 | 18 | 95% |
| Paper nonhits | 25 | 12 | 48% |

B hits:

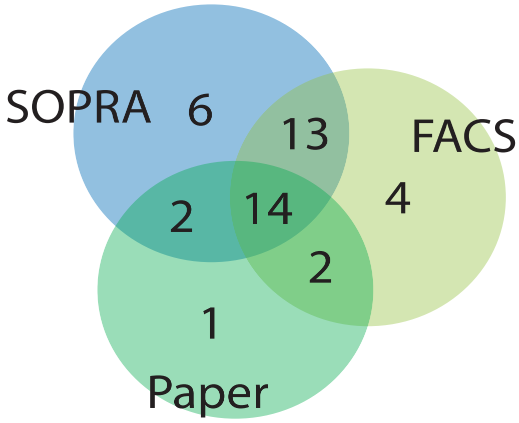


nonhits:

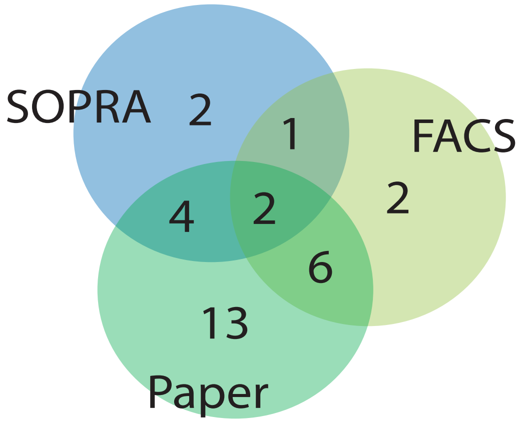
