## Supplementary Table 1 for "Single Object Profiles Regression Analysis (SOPRA): A novel method for analyzing high content cell-based screens": Supplementary Table 1.pdf

| GeneSymbol | Gene ID | # of siRNA tested |
|---|---|---|
| ABI1 | 10006 | 2 |
| ABI2 | 10152 | 2 |
| ACTR3 | 10096 | 1 |
| ADAM17 | 6868 | 1 |
| AKAP6 | 9472 | 1 |
| ALDH1A2 | 8854 | 1 |
| ANP32A | 8125 | 2 |
| AP1B1 | 162 | 2 |
| APAF1 | 317 | 2 |
| ARF4 | 378 | 2 |
| ARFGAP1 | 55738 | 2 |
| ATF2 | 1386 | 2 |
| ATG3 | 64422 | 2 |
| ATG4D | 84971 | 1 |
| ATG5 | 9474 | 2 |
| ATG9A | 79065 | 1 |
| ATR | 545 | 2 |
| AVEN | 57099 | 1 |
| AXIN2 | 8313 | 1 |
| BAG1 | 578 | 2 |
| BARD1 | 580 | 1 |
| BAT1 | 7919 | 2 |
| BCAT1 | 586 | 2 |
| BCL10 | 8915 | 1 |
| BCL2 | 596 | 2 |
| BCL2L1 | 598 | 2 |
| BCL2L11 | 10018 | 2 |
| BCL3 | 602 | 1 |
| BCR | 613 | 1 |
| BET1 | 10282 | 2 |
| BET1L | 51272 | 2 |
| BID | 637 | 2 |
| BIRC2 | 329 | 2 |
| BIRC3 | 330 | 1 |
| BIRC4 | 331 | 2 |
| BIRC5 | 332 | 2 |
| BMF | 90427 | 2 |
| BNIP1 | 662 | 1 |
| CALR | 811 | 1 |
| CAPN1 | 823 | 1 |
| CARD10 | 29775 | 1 |
| CASP1 | 834 | 2 |
| CASP14 | 23581 | 2 |

Hits

Non-Hits

| | | |
|---|---|---|
| CASP2 | 835 | 1 |
| CASP3 | 836 | 2 |
| CASP5 | 838 | 2 |
| CASP7 | 840 | 1 |
| CAV1 | 857 | 2 |
| CBL | 867 | 2 |
| CCNE1 | 898 | 1 |
| CCT2 | 10576 | 1 |
| CD40LG | 959 | 1 |
| CD44 | 960 | 1 |
| CD69 | 969 | 1 |
| CDC2 | 983 | 2 |
| CHEK1 | 1111 | 1 |
| CLK1 | 1195 | 2 |
| CLK2 | 1196 | 2 |
| CNN3 | 1266 | 1 |
| COPB | 1315 | 1 |
| CREB1 | 1385 | 2 |
| CARHSP1 | 23589 | 2 |
| CSK | 1445 | 2 |
| CTNNA1 | 1495 | 2 |
| DDX1 | 1653 | 1 |
| EIF3S4 | 8666 | 2 |
| FTL | 2512 | 1 |
| GABARAPL1 | 23710 | 1 |
| HNRPH1 | 3187 | 2 |
| HNRPK | 3190 | 1 |
| HNRPM | 4670 | 1 |
| IK | 3550 | 1 |
| INCENP | 3619 | 2 |
| KIF20A | 10112 | 1 |
| KIF3A | 11127 | 1 |
| KIF5B | 3799 | 1 |
| LIMK1 | 3984 | 2 |
| MCL1 | 4170 | 1 |
| MYC | 4609 | 2 |
| NCL | 4691 | 3 |
| NEK7 | 140609 | 2 |
| NXF1 | 10482 | 2 |
| PHB | 5245 | 2 |
| PICK1 | 9463 | 2 |
| PLCG2 | 5336 | 1 |
| PLK1 | 5347 | 2 |
| PRDX3 | 10935 | 2 |
| PRKD3 | 23683 | 1 |
| PRPF8 | 10594 | 2 |
| PSMA3 | 5684 | 2 |
| PSMA7 | 5688 | 1 |

| | | |
|---|---|---|
| PSMB1 | 5689 | 2 |
| PSMC3 | 5702 | 1 |
| PSMC4 | 5704 | 1 |
| PSMC5 | 5705 | 2 |
| PSMD7 | 5713 | 1 |
| RBBP4 | 5928 | 1 |
| ROCK1 | 6093 | 1 |
| RPLP0 | 6175 | 2 |
| SERPINB9 | 5272 | 2 |
| SF3A1 | 10291 | 2 |
| SNRPA1 | 6627 | 1 |
| TGFBR1 | 7046 | 1 |
| TNFRSF21 | 27242 | 1 |
| TSG101 | 7251 | 1 |
| WEE1 | 7465 | 2 |
| YWHAZ | 7534 | 2 |
