## Supplementary Table 2 for "Single Object Profiles Regression Analysis (SOPRA): A novel method for analyzing high content cell-based screens": Supplementary Table 2.pdf

| | | Wells | | | RNAs // Treatment | | | Hit // Nonhit | | | Benchmark | | | Images/FACS |
|---|---|---|---|---|---|---|---|---|---|---|---|---|---|---|
| | | Heatmap Group | | | Heatmap Group | | | Method | | | Method | | | |
| | ***Gene Names / Treatments*** | **1** | **2** | **3** | **1** | **2** | **3** | SOPRA | FACS | Paper | SOPRA | FACS | Paper | |
| 1 | *MYC* | 0 | 4 | 0 | 0 | 2 | 0 | hit | hit | hit | 1 | 1 | 1 | v |
| 2 | *CSK* | 0 | 4 | 0 | 0 | 2 | 0 | hit | hit | hit | 1 | 1 | 1 | v |
| 3 | *YWHAZ* | 0 | 2 | 0 | 0 | 1 | 0 | hit | | | | | | |
| 4 | *CCT2* | 0 | 2 | 0 | 0 | 1 | 0 | hit | | | | | | |
| 5 | *ABI2* | 0 | 2 | 0 | 0 | 1 | 0 | hit | | | | | | |
| 6 | *CDC2* | 0 | 4 | 0 | 0 | 2 | 0 | hit | hit | hit | 1 | 1 | 1 | v |
| 7 | *BCL2L11* | 0 | 3 | 0 | 0 | 2 | 0 | hit | | | | | | |
| 8 | *BET1L* | 0 | 4 | 0 | 0 | 2 | 0 | hit | | | | | | |
| 9 | *CAPN1* | 0 | 2 | 0 | 0 | 1 | 0 | hit | hit | nonhit | 1 | 1 | | v |
| 10 | *CLK1* | 0 | 3 | 0 | 0 | 2 | 0 | hit | | | | | | |
| 11 | *TNFRSF21* | 0 | 2 | 0 | 0 | 1 | 0 | hit | hit | hit | 1 | 1 | 1 | |
| 12 | *TGFBR1* | 0 | 2 | 0 | 0 | 1 | 0 | hit | hit | hit | 1 | 1 | 1 | |
| 13 | *ATR* | 0 | 2 | 0 | 0 | 1 | 0 | hit | hit | nonhit | 1 | 1 | | v |
| 14 | *KIF20A* | 0 | 2 | 0 | 0 | 1 | 0 | hit | hit | hit | 1 | 1 | 1 | |
| 15 | *PSMA7* | 0 | 2 | 0 | 0 | 1 | 0 | hit | hit | hit | 1 | 1 | 1 | |
| 16 | *ANP32A* | 0 | 2 | 0 | 0 | 1 | 0 | hit | nonhit | nonhit | | 1 | 1 | v |
| 17 | *CD69* | 0 | 2 | 0 | 0 | 1 | 0 | hit | nonhit | nonhit | | 1 | 1 | v |
| 18 | *BAT1* | 0 | 2 | 0 | 0 | 1 | 0 | hit | nonhit | hit | 1 | | 1 | v |
| 19 | *BCR* | 0 | 2 | 0 | 0 | 1 | 0 | hit | nonhit | nonhit | | 1 | 1 | v |
| 20 | *BARD1* | 0 | 2 | 0 | 0 | 1 | 0 | hit | | | | | | |
| 21 | *DDX1* | 0 | 2 | 0 | 0 | 1 | 0 | hit | | | | | | |
| 22 | *INCENP* | 0 | 2 | 0 | 0 | 1 | 0 | hit | | | | | | |
| 23 | *PSMA3* | 0 | 2 | 0 | 0 | 1 | 0 | hit | | | | | | |
| 24 | *CLK2* | 0 | 3 | 0 | 0 | 2 | 0 | hit | hit | nonhit | 1 | 1 | | |
| 25 | *CASP7* | 0 | 2 | 0 | 0 | 1 | 0 | hit | | | | | | |
| 26 | *NXF1* | 0 | 2 | 0 | 0 | 2 | 0 | hit | | | | | | |
| 27 | *AXIN2* | 0 | 1 | 0 | 0 | 1 | 0 | hit | | | | | | |
| 28 | *CAV1* | 0 | 2 | 0 | 0 | 2 | 0 | hit | | | | | | |
| 29 | *PICK1* | 0 | 2 | 0 | 0 | 1 | 0 | hit | hit | hit | 1 | 1 | 1 | |
| 30 | *BIRC2* | 0 | 2 | 0 | 0 | 2 | 0 | hit | | | | | | |
| 31 | *BET1* | 0 | 2 | 0 | 0 | 1 | 0 | hit | nonhit | nonhit | | 1 | 1 | |
| 32 | *ATG9A* | 0 | 1 | 0 | 0 | 1 | 0 | hit | | | | | | |
| 33 | *CD40LG* | 0 | 1 | 0 | 0 | 1 | 0 | hit | hit | nonhit | 1 | 1 | | |
| 34 | *ARF4* | 0 | 1 | 0 | 0 | 1 | 0 | hit | hit | hit | 1 | 1 | 1 | |
| 35 | *PSMB1* | 0 | 3 | 0 | 0 | 2 | 0 | hit | | | | | | |
| 36 | *BAG1* | 0 | 1 | 0 | 0 | 1 | 0 | hit | nonhit | nonhit | | 1 | 1 | |
| 37 | *CASP3* | 0 | 4 | 0 | 0 | 2 | 0 | hit | | | | | | |
| 38 | *MCL1* | 0 | 1 | 0 | 0 | 1 | 0 | hit | | | | | | |
| 39 | *AKAP6* | 0 | 2 | 0 | 0 | 1 | 0 | hit | | | | | | |
| 40 | *ATF2* | 0 | 2 | 0 | 0 | 1 | 0 | hit | | | | | | |
| 41 | *PRKD3* | 0 | 2 | 0 | 0 | 1 | 0 | hit | | | | | | |
| 42 | *PRDX3* | 0 | 1 | 0 | 0 | 1 | 0 | hit | hit | hit | 1 | 1 | 1 | |
| 43 | *A2* | 28 | 0 | 0 | 0 | 1 | 0 | hit | | | | | | |
| 44 | *A1* | 27 | 0 | 0 | 0 | 1 | 0 | hit | | | | | | |
| 45 | *SNRPA1* | 2 | 0 | 0 | 0 | 1 | 0 | hit | | | | | | |
| 46 | *Allstars* | 2 | 0 | 0 | 0 | 1 | 0 | hit | | | | | | |
| 47 | *COPB* | 1 | 0 | 0 | 0 | 1 | 0 | hit | hit | hit | 1 | 1 | 1 | |
| 48 | *AP1B1* | 4 | 0 | 0 | 2 | 0 | 0 | hit | | | | | | |
| 49 | *CTNNA1* | 2 | 0 | 0 | 1 | 0 | 0 | hit | hit | nonhit | 1 | 1 | | |
| 50 | *NCL* | 4 | 0 | 0 | 2 | 0 | 0 | hit | | | | | | |
| 51 | *CARD10* | 1 | 0 | 0 | 1 | 0 | 0 | hit | | | | | | |
| 52 | *FTL* | 1 | 0 | 0 | 1 | 0 | 0 | hit | | | | | | |
| 53 | *CD44* | 1 | 0 | 0 | 1 | 0 | 0 | hit | | | | | | |
| 54 | *MockI* | 3 | 0 | 0 | 2 | 0 | 0 | hit | | | | | | |
| 55 | *PSMD7* | 2 | 0 | 0 | 1 | 0 | 0 | hit | | | | | | |
| 56 | *ATG5* | 2 | 0 | 0 | 1 | 0 | 0 | hit | hit | nonhit | 1 | 1 | | |
| 57 | *CRHSP24* | 3 | 0 | 0 | 2 | 0 | 0 | hit | | | | | | |
| 58 | *TSG101* | 2 | 0 | 0 | 1 | 0 | 0 | hit | | | | | | |
| 59 | *CREB1* | 2 | 0 | 0 | 1 | 0 | 0 | hit | hit | nonhit | 1 | 1 | | |
| 60 | *ACTR3* | 2 | 0 | 0 | 1 | 0 | 0 | hit | | | | | | |
| 61 | *CHEK1* | 2 | 0 | 0 | 1 | 0 | 0 | hit | | | | | | |
| 62 | *CCNE1* | 2 | 0 | 0 | 1 | 0 | 0 | hit | | | | | | |
| 63 | *BCL10* | 1 | 0 | 0 | 1 | 0 | 0 | hit | | | | | | |
| 64 | *BIRC5* | 3 | 0 | 0 | 2 | 0 | 0 | hit | | | | | | |
| 65 | *HNRPK* | 2 | 0 | 0 | 1 | 0 | 0 | hit | | | | | | |
| 66 | *BIRC4* | 3 | 0 | 0 | 2 | 0 | 0 | hit | | | | | | |
| 67 | *ARFGAP1* | 2 | 0 | 0 | 1 | 0 | 0 | hit | hit | nonhit | 1 | 1 | | |
| 68 | *CASP14* | 3 | 0 | 0 | 2 | 0 | 0 | hit | | | | | | |
| 69 | *EIF3S4* | 3 | 0 | 0 | 2 | 0 | 0 | hit | | | | | | |
| 70 | *KIF5B* | 1 | 0 | 0 | 1 | 0 | 0 | hit | | | | | | |
| 71 | *Luci* | 4 | 0 | 0 | 2 | 0 | 0 | hit | | | | | | |
| 72 | *KIF3A* | 2 | 0 | 0 | 1 | 0 | 0 | hit | | | | | | |
| 73 | *CBL* | 3 | 0 | 0 | 2 | 0 | 0 | hit | | | | | | |
| 74 | *AVEN* | 1 | 0 | 0 | 1 | 0 | 0 | hit | | | | | | |
| 75 | *RPLP0* | 4 | 0 | 0 | 2 | 0 | 0 | hit | | | | | | |
| 76 | *WEE1* | 1 | 0 | 0 | 1 | 0 | 0 | hit | | | | | | |
| 77 | *RBBP4* | 1 | 0 | 0 | 1 | 0 | 0 | hit | | | | | | |
| 78 | *PLCG2* | 1 | 0 | 0 | 1 | 0 | 0 | hit | | | | | | |
| 79 | *A3* | 2 | 26 | 0 | 1 | 0 | 0 | hit | | | | | | |
| 80 | *A4* | 7 | 21 | 0 | 1 | 0 | 0 | hit | | | | | | |
| 81 | *LIMK1* | 1 | 1 | 0 | 1 | 1 | 0 | weak hit** | hit | nonhit | 1 | 1 | | |
| 82 | *BCL2L1* | 2 | 2 | 0 | 1 | 1 | 0 | weak hit** | | | | | | |
| 83 | *ABI1* | 2 | 1 | 0 | 1 | 1 | 0 | weak hit** | hit | nonhit | 1 | 1 | | |
| 84 | *ATG3* | 2 | 2 | 0 | 1 | 1 | 0 | weak hit** | hit | nonhit | 1 | 1 | | |
| 85 | *SF3A1* | 3 | 1 | 0 | 2 | 1 | 0 | medium hit* | | | | | | |
| 86 | *CASP1* | 1 | 3 | 0 | 2 | 1 | 0 | medium hit* | hit | nonhit | 1 | 1 | | |
| 87 | *APAF1* | 1 | 3 | 0 | 1 | 2 | 0 | medium hit* | nonhit | nonhit | | 1 | 1 | |
| 88 | *BCL2* | 2 | 2 | 0 | 1 | 1 | 0 | weak hit** | | | | | | |
| 89 | *BNIP1* | 1 | 1 | 0 | 1 | 1 | 0 | weak hit** | | | | | | |
| 90 | *PHB* | 2 | 2 | 0 | 1 | 1 | 0 | weak hit** | hit | hit | 1 | 1 | 1 | |
| 91 | *HNRPM* | 1 | 1 | 0 | 1 | 1 | 0 | weak hit** | nonhit | hit | 1 | | 1 | |
| 92 | *BCAT1* | 3 | 1 | 0 | 2 | 1 | 0 | medium hit* | | | | | | |
| 93 | *PLK1* | 1 | 1 | 0 | 1 | 1 | 0 | weak hit** | | | | | | |
| 94 | *BMF* | 1 | 2 | 0 | 1 | 1 | 0 | weak hit** | | | | | | |
| 95 | *BID* | 1 | 1 | 0 | 1 | 1 | 0 | weak hit** | | | | | | |
| 96 | *HNRPH1* | 3 | 1 | 0 | 2 | 1 | 0 | weak hit** | | | | | | |
| 97 | *PSMC5* | 2 | 1 | 0 | 2 | 1 | 0 | medium hit* | hit | hit | 1 | 1 | 1 | |
| 98 | *CASP5* | 1 | 1 | 0 | 1 | 1 | 0 | weak hit** | hit | nonhit | 1 | 1 | | |
| 99 | *SERPINB9* | 1 | 1 | 0 | 1 | 1 | 0 | weak hit** | hit | hit | 1 | 1 | 1 | |
| 100 | *N2* | 0 | 0 | 28 | 0 | 0 | 1 | hit | | | | | | |
| 101 | *N1* | 0 | 0 | 28 | 0 | 0 | 1 | hit | | | | | | |
| 102 | *N4* | 0 | 0 | 28 | 0 | 0 | 1 | hit | | | | | | |
| 103 | *N3* | 1 | 0 | 27 | 1 | 0 | 1 | medium hit* | | | | | | |
| 104 | CALR | 0 | 0 | 0 | 0 | 0 | 0 | nonhit | nonhit | nonhit | 1 | 1 | 1 | |
| 105 | CASP2 | 0 | 0 | 0 | 0 | 0 | 0 | nonhit | nonhit | hit | 1 | 1 | | |
| 106 | ALDH1A2 | 0 | 0 | 0 | 0 | 0 | 0 | nonhit | nonhit | nonhit | 1 | 1 | 1 | |
| 107 | BCL3 | 0 | 0 | 0 | 0 | 0 | 0 | nonhit | hit | nonhit | 1 | | 1 | |
| 108 | BIRC3 | 0 | 0 | 0 | 0 | 0 | 0 | nonhit | hit | nonhit | 1 | | 1 | |
| 109 | ROCK1 | 0 | 0 | 0 | 0 | 0 | 0 | nonhit | hit | hit | | 1 | 1 | |
| 110 | CNN3 | 0 | 0 | 0 | 0 | 0 | 0 | nonhit | hit | nonhit | 1 | | 1 | |
| 111 | ADAM17 | 0 | 0 | 0 | 0 | 0 | 0 | nonhit | hit | nonhit | 1 | | 1 | |
| 112 | YWHA2 | 0 | 0 | 0 | 0 | 0 | 0 | nonhit | hit | hit | | 1 | 1 | |
| | | | | | | | | | | | 36 | 38 | 30 | |

** 2 siRNAs in different heatmap groups

* 2 siRNAs in different heatmap groups with a tendency to one.
