## Supplementary File 1 for "Single Object Profiles Regression Analysis (SOPRA): A novel method for analyzing high content cell-based screens": SOPRA-Project_Description-modified-by-KPP-version-16.pdf

Authors: André P. Maeurer, Klaus-Peter Pleissner

Version 0.15
Date: 2021/01/10

#### 1 Introduction

The analysis of high-content data, such as cell-based feature data derived from image analysis from RNAi screens or FACS data poses an important research field in the genomic area.

The Single Object Profiles Regression Analysis (SOPRA) (Fig.1) workflow enables researchers to identify cell populations with statistically significant changes of normalized frequency distribution profiles (histograms) of measured cellular features based on the regression analysis. The regression-based approach was performed using maSigPro from Bioconductor, an R-package originally applied for time-course microarray analysis. We defined a regression model where the dependent variable was the bin-wise normalized frequency distribution profile and the independent variable was the measured cellular feature within predefined binning intervals. Our experimental design was based on the single series time-course approach of maSigPro. Shortly, maSigPro follows a two steps regression strategy to find genes with significant temporal expression changes and significant differences between experimental groups. The method defines a general regression model for the data where the experimental groups are identified by dummy variables. The procedure first adjusts this global model by the least-squared technique to identify differentially expressed genes and selects significant genes applying false discovery rate control procedures. Secondly, stepwise regression is applied as a variable selection strategy to study differences between experimental groups and to find statistically significant different profiles. The coefficients obtained in this second regression model will be useful to cluster together significant genes with similar expression patterns and to visualize the results. These explanations are given in the maSigPro user’s guide.
The workflow can easily be run on analyzed images or FACS data from thousands of cell populations.

#### 2 Description of software

The SOPRA software workflow (Fig.2) is realized as a shiny application with an user interface UI.R and application SERVER.R and mainly consists of four parts, (SOPRA 1 of 4) preprocessing, (SOPRA 2 of 4) data gathering and normalization, (SOPRA 3 of 4) identification of significantly changed cell populations and clustering and (SOPRA 4 of 4) conversion of these significant findings into genes with their cluster membership.

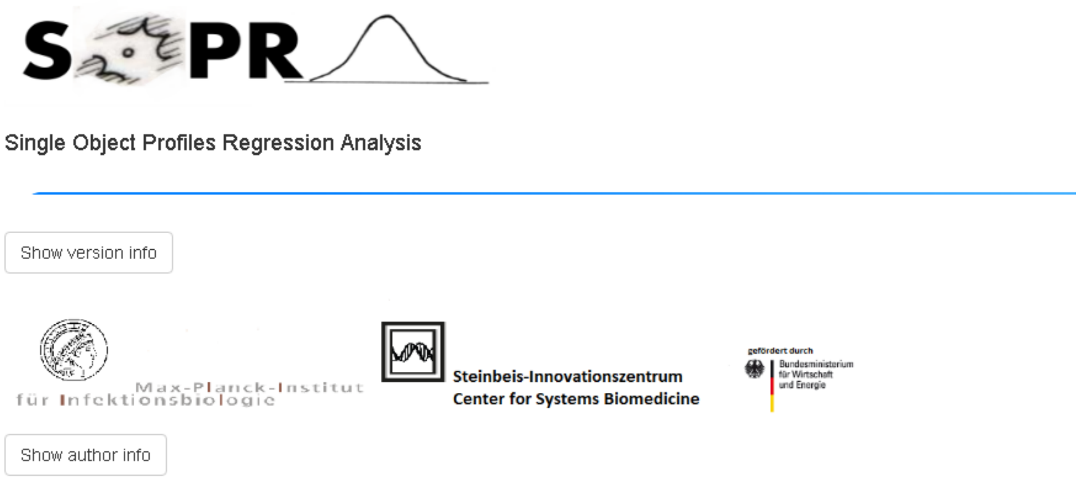


Fig.1 Header of the SOPRA User Interface

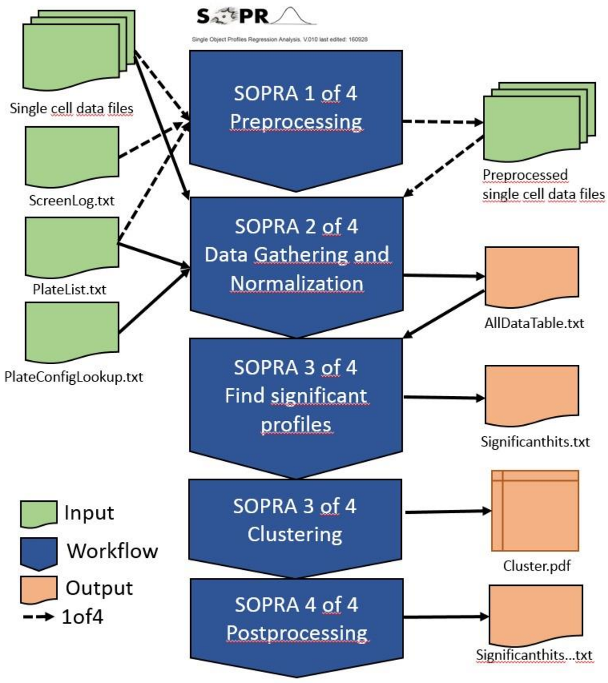


Fig.2 SOPRA workflow

#### 2.1 Preprocessing

Due to technical or other reasons some wells can contain faulty information. Therefore, these wells must be excluded from further analysis. SOPRA 1 of 4 (Preprocessing) performs a preprocessing step, in which all features of the indicated incorrect wells are flagged by NA. The specified wells are listed in the manually-created 'ScreenLog' file (for instance: ScreenLog-384.txt).

#### 2.2 Data Gathering and Normalization

SOPRA 2 of 4 (Data Gathering and Normalization) consists of several steps leading to normalized histograms, in which the original data values within a given interval (bin) are replaced normalized density distribution (histogram).

First, all data is read according to the file 'PlateList.txt' and annotates all objects (single cell data) with the data from the file 'PlateConf_Lookup.txt'. INF-values are flagged and commas replaced with dotted decimal notation.

Several image analysis (such as SCAN^R) or FACS software programs offer the opportunity to flag objects if they are not in a specific gate. In a next (optional) step we use the flagging information stored in the data files and the 'PlateConf_LookUp.txt' and 'PlateList.txt' file to filter these objects. The 'PlateConf_LookUp.txt' file contains the column 'Filter' with the active filter combinations (e.g. F1 or F1&F2), the 'PlateList.txt' contains the names of the filters (e.g. the column for the Area gating has been given the name 'G_Area41726561' by SCAN^R). After this step, the data is exported as 'Data_rawdata_PlatexReplicatey.txt'.

Next, the quantiles and median raw values of the control wells are calculated and an object-wise median of control (moc) normalization for each plate (called: plate-wise) is performed. After this step, the data is exported as 'Data_nzdata_PlatexReplicatey.txt'.

Then, the binning-(interval)-axis is calculated for the complete data set.

Next, QC blots are exported for the normalized data.

After that, histograms based on the binning-axis are calculated for all sample and control wells, resulting in a certain number of objects (counts) for each bin of a well. For all sample and control wells, the counts of each bin are divided by the median count of the respective bin of the control wells to perform a 'bin-wise' normalization, leading to a log2 normalized frequency distribution (histogram) for all samples and controls. The larger the deviation of a sample from the reference (control) distribution, the more likely the test distribution is statistically significant.

Finally, the data is saved in the file 'AllDataTable.txt'.

#### 2.3 Identify statistically significant frequency distribution profiles

SOPRA 3 of 4 (Find Statistically Significant Profiles) identifies log2 normalized frequency distributions that are significantly altered compared to the control distributions, using the Bioconductor R-package 'maSigPro' in the single series time-course approach (for further details see the maSigPro user's guide).

#### 2.4 Post-processing

SOPRA 4 of 4 (siRNA to Gene Post Processing) converts the siRNA hit list into a list containing the gene symbols given in the PlateConfLookUp.txt table, the cluster membership, and the frequency in the corresponding cluster. Because post-processing makes always sense it is the final step of SOPRA 3 of 4 software part.

#### 3 Input Sources

The single cell data files and the descriptive files for 384-well plates from our cell cycle screen can be downloaded from MPIIB cloud server MPIIB cloud server .

The single cell data (Fig. 3) is stored in tab-delimited files. Each row contains the data for one cell (object). Each file must contain an ID and the corresponding well and at least one measured feature for each object. Therefore, the objects do not have to be in a chronological order. Optionally, if you want to use the gating filter, it can also contain several gating filters (e.g. ‘G_Area41726561’; ‘1’ = object inside gate, ‘0’ = object outside gate).

We programmed an image analysis assay for the ‘Scan^R’ software (Olympus) to generate single object data. This tab-delimited data files can contain upto 900000 raws (objects) for one replicate of one plate and can be opened by EXCEL. The file structure looks as follows:

| Object ID | Total Intensity DAPI | Mean Intensity DAPI | Area | Well | G_Area41726561 | G_MeanDAPI4D65616E44415049 | G_TotalDAPI546F74616C44415049 |
|---|---|---|---|---|---|---|---|
| 0 | 320989 | 840.28534 | 382 | 1 | 1 | 1 | 1 |
| 1 | 691633 | 1000.9161 | 691 | 1 | 1 | 1 | 1 |
| 2 | 317899 | 572.79102 | 555 | 1 | 1 | 1 | 1 |
| 3 | 820896 | 987.84113 | 831 | 1 | 1 | 1 | 1 |
| 4 | 352608 | 993.26196 | 355 | 1 | 1 | 1 | 1 |
| 5 | 381844 | 1243.7915 | 307 | 1 | 1 | 1 | 1 |
| 6 | 892706.94 | 739.60809 | 1207 | 1 | 1 | 1 | 1 |
| 7 | 361899 | 1052.032 | 344 | 1 | 1 | 1 | 1 |
| 8 | 729447 | 1146.9292 | 636 | 1 | 1 | 1 | 1 |

Fig.3 File structure for the data files (some columns are

blanked out).

The following tab-delimited descriptive files are needed for

the workflow.

- PlateList.txt

The file ‘PlateList.txt‘ (Fig. 4) contains all necessary information about the screening experiment. For each feature to analyze a respective file has to be generated. It must contain the *well range* of the file, the *file name*, the *plate* and *replicate number* of the plate, the *control* and the *feature* to be analyzed (connected to a feature column of the single cell data files).
Optionally, you can use up to three cell gating filters (connected to a gating column of the single cell data files)

| Platename | WellRange | ObjectType | Filename | Plate | Replicate | Control | Feature | Filter1 | Filter2 | Filter3 |
|---|---|---|---|---|---|---|---|---|---|---|
| Plate001_R1 | W1-384 | Main | Plate001_R1_W1-384_Main.txt | 1 | 1 | NeutralControl1 | Area | G_Nuclei4E75636C6569 | G_Area41726561 | NA |
| Plate001_R2 | W1-384 | Main | Plate001_R2_W1-384_Main.txt | 1 | 2 | NeutralControl1 | Area | G_Nuclei4E75636C6569 | G_Area41726561 | NA |
| Plate001_R3 | W1-384 | Main | Plate001_R3_W1-384_Main.txt | 1 | 3 | NeutralControl1 | Area | G_Nuclei4E75636C6569 | G_Area41726561 | NA |
| Plate001_R4 | W1-384 | Main | Plate001_R4_W1-384_Main.txt | 1 | 4 | NeutralControl1 | Area | G_Nuclei4E75636C6569 | G_Area41726561 | NA |
| Plate002_R1 | W1-384 | Main | Plate002_R1_W1-384_Main.txt | 2 | 1 | NeutralControl1 | Area | G_Nuclei4E75636C6569 | G_Area41726561 | NA |
| Plate002_R2 | W1-384 | Main | Plate002_R2_W1-384_Main.txt | 2 | 2 | NeutralControl1 | Area | G_Nuclei4E75636C6569 | G_Area41726561 | NA |
| Plate002_R3 | W1-384 | Main | Plate002_R3_W1-384_Main.txt | 2 | 3 | NeutralControl1 | Area | G_Nuclei4E75636C6569 | G_Area41726561 | NA |
| Plate002_R4 | W1-384 | Main | Plate002_R4_W1-384_Main.txt | 2 | 4 | NeutralControl1 | Area | G_Nuclei4E75636C6569 | G_Area41726561 | NA |

Fig. 4 File structure for the ‘PlateList.txt’ for a 384 well plate

Sometimes, due to the large amount of data generated during the single cell analysis, it is impossible to export all data in one file. Therefore, it is possible to export parts of a complete plate, such as 4 files containing data for 96 wells each, for a 384 well plate. The parts themselves have to be chronological, however the wells inside the files themselves (such as well 1-96) don’t have to be chronological.

For the generation of the manuscript data we used the 1*384 data structure version with Area as selected feature.

- PlateConfLookUp.txt

The file ‘PlateConfLookUp.txt’ (Fig. 5) has to be prepared once and contains all necessary library information, such as plate number, well annotation, well content and gene symbol.

Also the information on which wells should be used as controls (such as ‘NeutralControl1’) is derived from the column ‘Content’. Which control is used for which plate is stored in the ‘PlateList.txt’ in the column ‘Control’.

Additionally, in the ‘Processing’ column of the ‘PlateConfLookUp.txt’ the user can define whether certain wells should be removed from the analysis. This is important for wells that should generally not be part of the analysis, such as toxic controls or outer wells, and are the same across all replicates used. To remove wells that are different between the replicates use the ‘ScreenLog.txt’ file.

In the column ‘Filter’ predefined filters can be used (such as F1&F2 for Filter1 and Filter2) for the object gating option. The names of the filters are derived from the ‘PlateList.txt’ file and only objects that meet the gating criteria and are characterized in the single cell data file as ‘1’ in the respective Filter column are included in the analysis.

| Plate | Well_Annotation_1 | Well_Annotation_2 | Content | GeneSymbol | Processing | Filter |
|---|---|---|---|---|---|---|
| 1 | A01 | 1 | OuterWell | Mock | FALSE | F1&F2 |
| ... | | | | | | |
| 1 | B09 | 33 | Sample | PHB_siRNA-43 | TRUE | F1&F2 |
| 1 | B10 | 34 | Sample | BAG1_siRNA-57 | TRUE | F1&F2 |
| 1 | B11 | 35 | Sample | BAG1_siRNA-57 | TRUE | F1&F2 |
| 1 | B12 | 36 | Sample | BCL10_siRNA-71 | TRUE | F1&F2 |
| 1 | B13 | 37 | Sample | BCL10_siRNA-71 | TRUE | F1&F2 |
| 1 | B14 | 38 | NeutralControl1 | Allstars | TRUE | F1&F2 |
| 1 | B15 | 39 | NeutralControl3 | MockI | TRUE | F1&F2 |
| 1 | B16 | 40 | InhibitorControl1 | A1 | TRUE | F1&F2 |
| 1 | B17 | 41 | InhibitorControl2 | A2 | TRUE | F1&F2 |
| 1 | B18 | 42 | InhibitorControl5 | N1 | TRUE | F1&F2 |
| 1 | B19 | 43 | InhibitorControl6 | N2 | TRUE | F1&F2 |
| 1 | B20 | 44 | InhibitorControl3 | A3 | TRUE | F1&F2 |
| 1 | B21 | 45 | InhibitorControl4 | A4 | TRUE | F1&F2 |
| 1 | B22 | 46 | InhibitorControl7 | N3 | TRUE | F1&F2 |
| 1 | B23 | 47 | InhibitorControl8 | N4 | TRUE | F1&F2 |
| 1 | B24 | 48 | ToxicControl | Plk 1-1µM | FALSE | F1&F2 |
| 1 | C01 | 49 | OuterWell | Mock | FALSE | F1&F2 |
| 1 | C02 | 50 | Sample | CRHSP24_siRNA-2 | TRUE | F1&F2 |
| ... | | | | | | |
| 2 | P22 | 382 | OuterWell | Mock | FALSE | F1&F2 |
| 2 | P23 | 383 | OuterWell | Mock | FALSE | F1&F2 |
| 2 | P24 | 384 | OuterWell | Mock | FALSE | F1&F2 |

Fig. 5 File structure for the ‘PlateConfLookUp.txt’

- ScreenLog.txt (optional)

The file ‘ScreenLog.txt’ (Fig. 6) is optional and contains information about wells that should be dismissed from the analysis but differ among the replicates of a plate. This might be the case if the experimenter notices an error in a certain well (such as a contamination) or certain wells do not meet quality criteria post screening (such as a minimum cell number).
The ‘ScreenLog.txt’ file is only needed if the SOPRA 1 of 4 (Preprocessing) option is used.

| Plate | Replicate | Well | WellNo | Flag | Comment | Barcode | Inducer |
|---|---|---|---|---|---|---|---|
| 1 | 1 | K8 | 248 | NA | no siRNA | Plate001_R1_W1-384_Main.txt | none |
| 1 | 1 | K9 | 249 | NA | no siRNA | Plate001_R1_W1-384_Main.txt | none |
| 1 | 2 | K8 | 248 | NA | no siRNA | Plate001_R2_W1-384_Main.txt | none |
| 1 | 2 | K9 | 249 | NA | no siRNA | Plate001_R2_W1-384_Main.txt | none |
| 1 | 3 | K8 | 248 | NA | no siRNA | Plate001_R3_W1-384_Main.txt | none |
| 1 | 3 | K9 | 249 | NA | no siRNA | Plate001_R3_W1-384_Main.txt | none |
| 1 | 4 | K8 | 248 | NA | no siRNA | Plate001_R4_W1-384_Main.txt | none |
| 1 | 4 | K9 | 249 | NA | no siRNA | Plate001_R4_W1-384_Main.txt | none |
| 2 | 1 | O12 | 348 | NA | no siRNA | Plate002_R1_W1-384_Main.txt | none |
| 2 | 1 | O13 | 349 | NA | no siRNA | Plate002_R1_W1-384_Main.txt | none |
| 2 | 2 | O12 | 348 | NA | no siRNA | Plate002_R2_W1-384_Main.txt | none |
| 2 | 2 | O13 | 349 | NA | no siRNA | Plate002_R2_W1-384_Main.txt | none |
| 2 | 3 | O12 | 348 | NA | no siRNA | Plate002_R3_W1-384_Main.txt | none |
| 2 | 3 | O13 | 349 | NA | no siRNA | Plate002_R3_W1-384_Main.txt | none |
| 2 | 4 | O12 | 348 | NA | no siRNA | Plate002_R4_W1-384_Main.txt | none |
| 2 | 4 | O13 | 349 | NA | no siRNA | Plate002_R4_W1-384_Main.txt | none |

Fig.6 File structure for the ‘ScreenLog.txt’

/!\ Filter options:

SOPRA offers the possibility to remove wells for all replicates from the analysis using either the column ‘Processing’ in the file ‘PlateConfLookUp.txt’, or for a defined replicate using the file ‘ScreenLog.txt’. Additionally, in wells that are analyzed individual objects can be dismissed from the analysis using the gating filter settings in the single cell data files and the files ‘PlateList.txt’ and ‘PlateConfLookUp.txt’.

### 4. Analysis

#### 4.1 Install R-Studio and R

First of all install R-Studio (https://www.rstudio.com/) and R (https://www.r-project.org/). We used R-Studio Version 1.3.1093 and R version 4.0.3 for Windows and RStudio-Server with Rstudio 1.1.463 and R-version 3.4.3 for Linux.

#### 4.2 Start the analysis

1. The software is realized as a shiny application with an user interface ui.R and application server.R within the R-Project SOPRA.Rproj
2. SOPRA shiny application (ui.R, server.R) and single cell data ( 96-well plates) for testing are accessible from GitHub https://github.com/kppleissner/SOPRA/. Download the entire repository and unzip it locally.
3. 384-well plates for SOPRA are available from MPIIB cloud server https://transfer.mpiib-berlin.mpg.de/s/AibR4AHLCR9xzDB?path=%2F
4. Launch R-Studio
5. Open the project SOPRA.Rproj in RStudio
6. After loading click on the button Run App
7. R-packages that are necesseray are installed automatically. Otherwise install these package manually.
8. Run through the user interface, set input/output folders for data, upload descriptive files and set corresponding parameters
9. Hit ‘Go Analysis’

#### 4.3 Data input and output, uploading Platelist.txt and PlateConfigLookUp.txt

The single cell data files to be read are determined (Select Input Folder) and the files ‘PlateList.txt and ‘PlateConfLookup.txt’ can be uploaded. Additionally you can define where to store the output files (Select Output Folder)-Fig. 7.

#### Folder selection

**Choose the input folder with the ScanR data files**

Select Input folder

**Following input folder was selected**

M:/SOPRAData/SOPRAdata-384/InData384

**Choose the output folder for resulting data**

Select or create Output folder

**Following output folder was selected**

M:/Area

#### File selection

**Select Platelist file with information which files should be processed**

Choose Platelist-file

**Following Platelist was choosen:**

[1] "M:/SOPRAData/SOPRAdata-384/Files384/PlateList-Area-384-with_NeutralC1.txt"

**Select PlateConfLookUp file with information on plate configuation**

Choose PlateConfLookUp-file

**Following PlateConfLookUp was choosen:**

[1] "M:/SOPRAData/SOPRAdata-384/Files384/PlateConf_LookUp.txt"

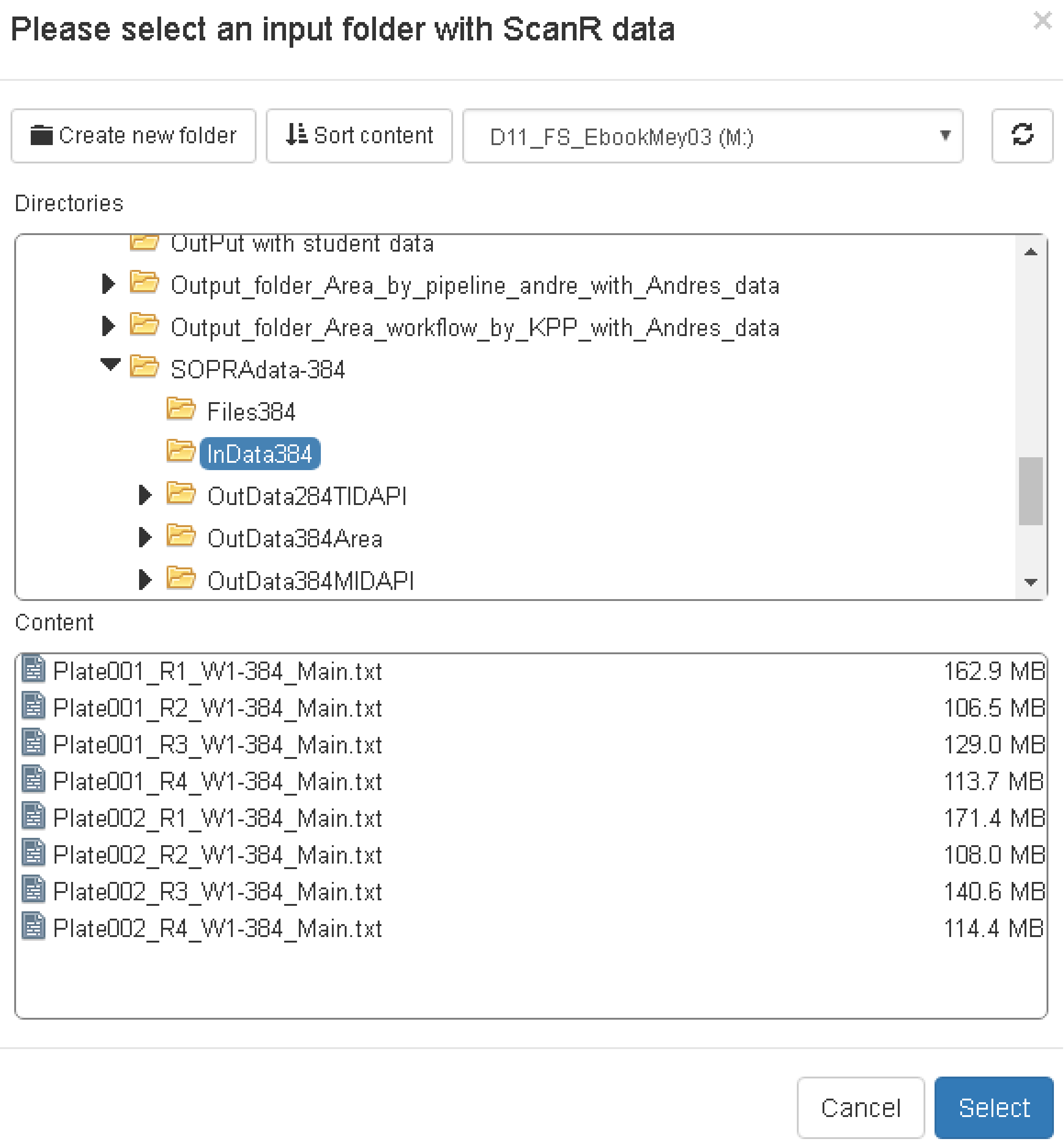
Please select an input folder with ScanR data
Create new folder
Sort content
D11_FS_EbookMey03 (M:)
Directories
OutPut with student data
Output_folder_Area_by_pipeline_andre_with_Andres_data
Output_folder_Area_workflow_by_KPP_with_Andres_data
SOPRAdata-384
Files384
InData384
OutData284TIDAPI
OutData384Area
OutData384MIDAPI
Content
Plate001_R1_W1-384_Main.txt 162.9 MB
Plate001_R2_W1-384_Main.txt 106.5 MB
Plate001_R3_W1-384_Main.txt 129.0 MB
Plate001_R4_W1-384_Main.txt 113.7 MB
Plate002_R1_W1-384_Main.txt 171.4 MB
Plate002_R2_W1-384_Main.txt 108.0 MB
Plate002_R3_W1-384_Main.txt 140.6 MB
Plate002_R4_W1-384_Main.txt 114.4 MB
Cancel
Select

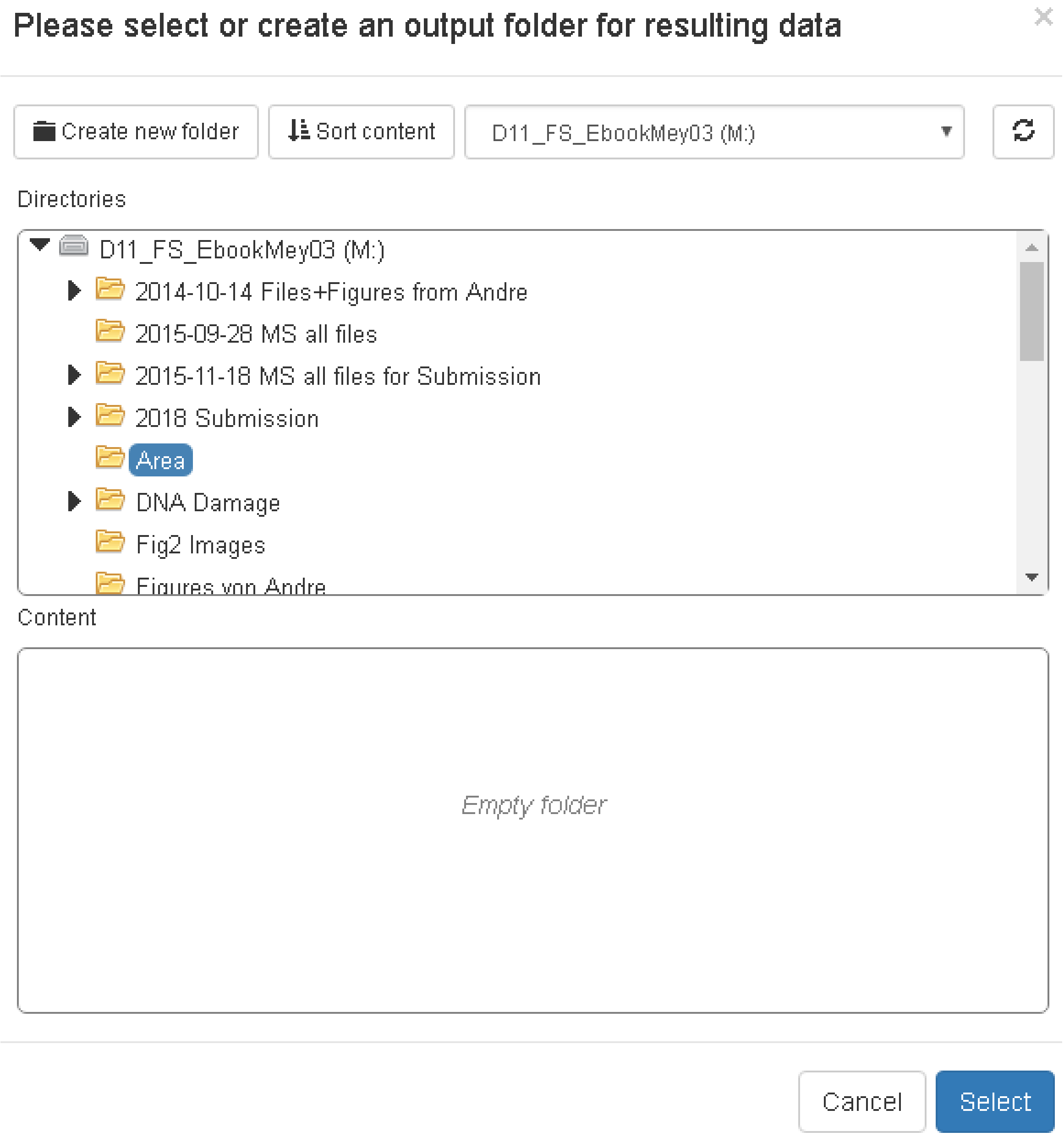
Please select or create an output folder for resulting data
Create new folder
Sort content
D11_FS_EbookMey03 (M:)
Directories
D11_FS_EbookMey03 (M:)
2014-10-14 Files+Figures from Andre
2015-09-28 MS all files
2015-11-18 MS all files for Submission
2018 Submission
Area
DNA Damage
Fig2 Images
Content
Empty folder
Cancel
Select

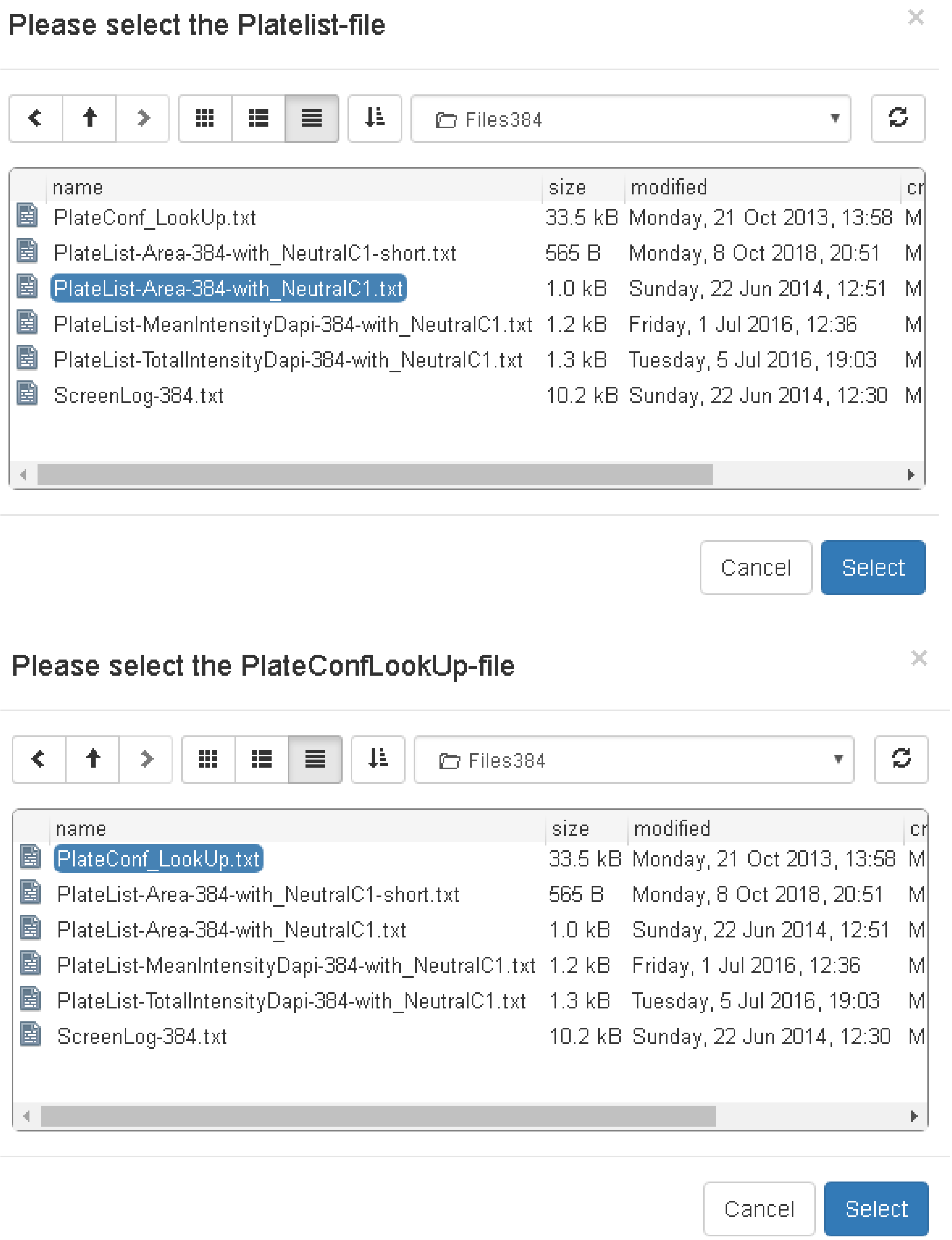


Fig. 7 File upload menu

#### 4.4 SOPRA 1 of 4 (Preprocessing)

The preprocessing is an optional step (Fig. 8) for which the ‘ScreenLog.txt’ has to be prepared as described above. It can be used for flagging wells that should be removed from the analysis but differ among the replicates of a plate. If the checkbox is activated, the SOPRA workflow generates the subfolder Preprocessing_Results in the dedicated output folder to store the preprocessed files.

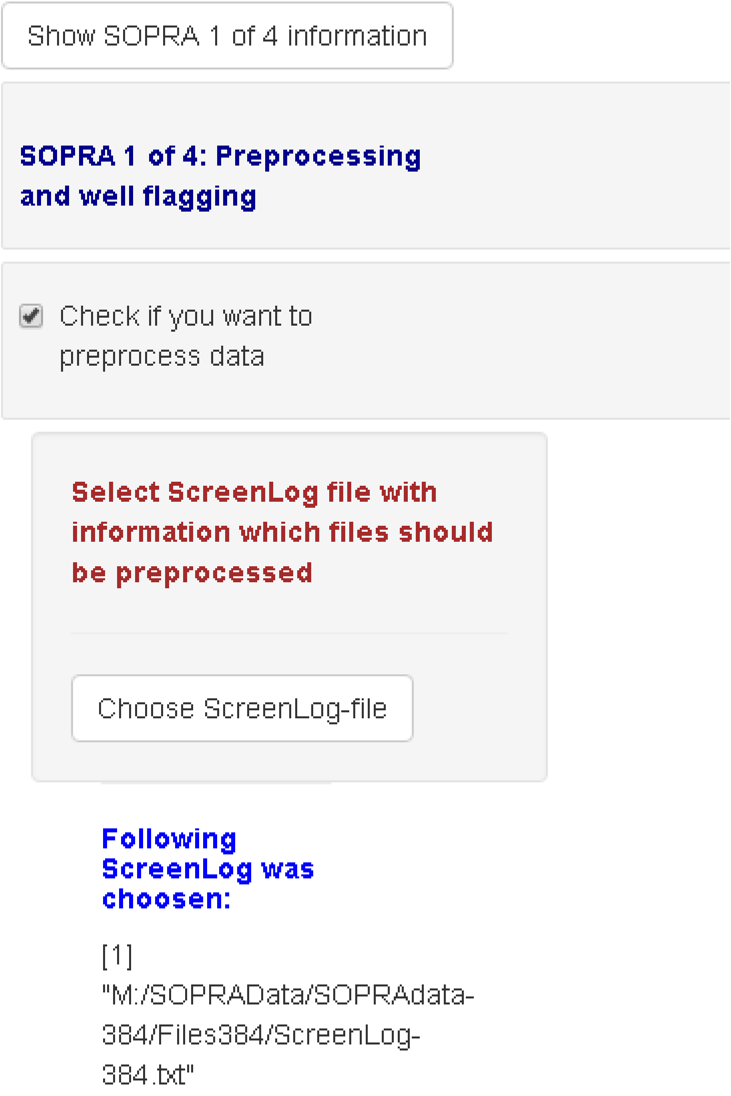

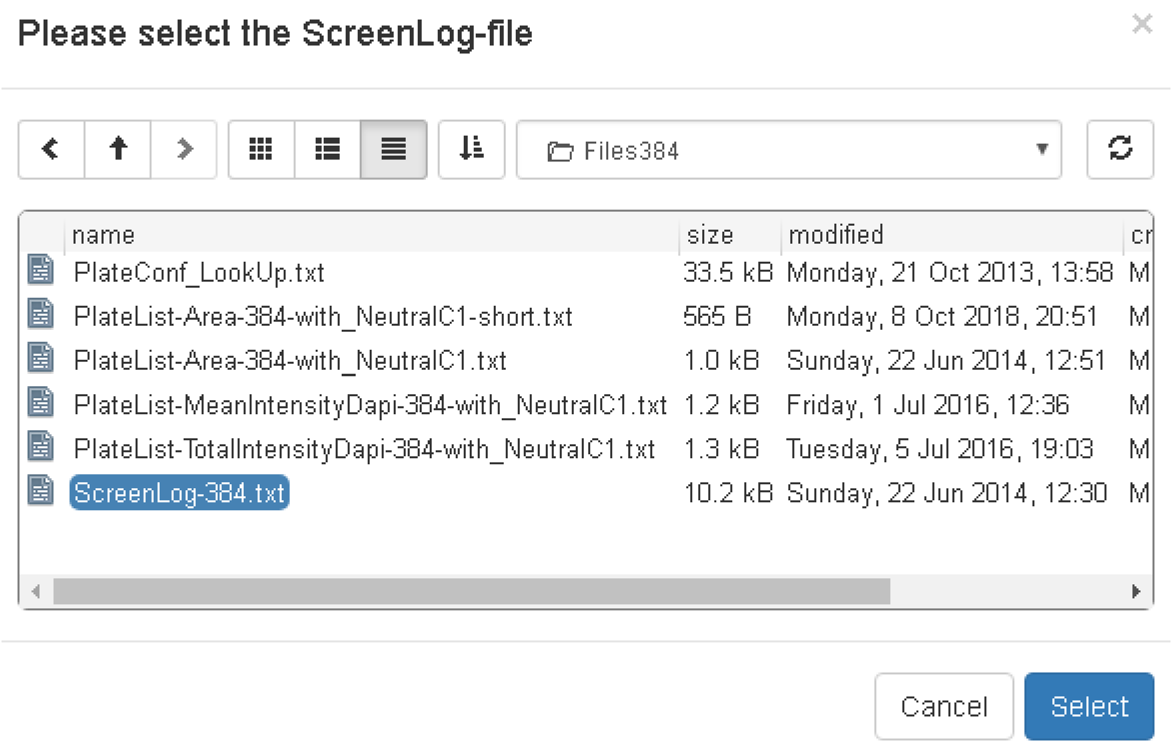


Fig. 8 Menu for SOPRA 1 of 4 (Preprocessing)

#### 4.5 SOPRA 2 of 4 (Data Gathering and Normalization)

For the data gathering and normalization step (Fig. 9), the number of bins for the generation of histograms has to be defined. Additionally the plate format and the left and right quantiles have to be set. The quantile settings offer the option to remove objects from the analysis steps, to avoid that outliers lead to mostly empty bins, which might result in abnormal results. Additionally, objects that have been predefined in the image analysis software to be outside of a certain gating range (see single cell data files) can be removed from further analysis by setting the gating filter options to filter ‘1-3’ (see ‘PlateList.txt’) or to the predefined filter settings (see ‘PlateConfLookUp.txt’). Further, it is possible to define whether the intermediate data files, the QC blots and a file containing the binning parameters should be exported. The external binning file is mandatory if SOPRA 3 of 4 (Find Significant Profiles) and the following steps should be executed without performing SOPRA 2 of 4 (Data Gathering and Normalization) again. All data are exported into the file “AllDataTable.txt” in the output folder.

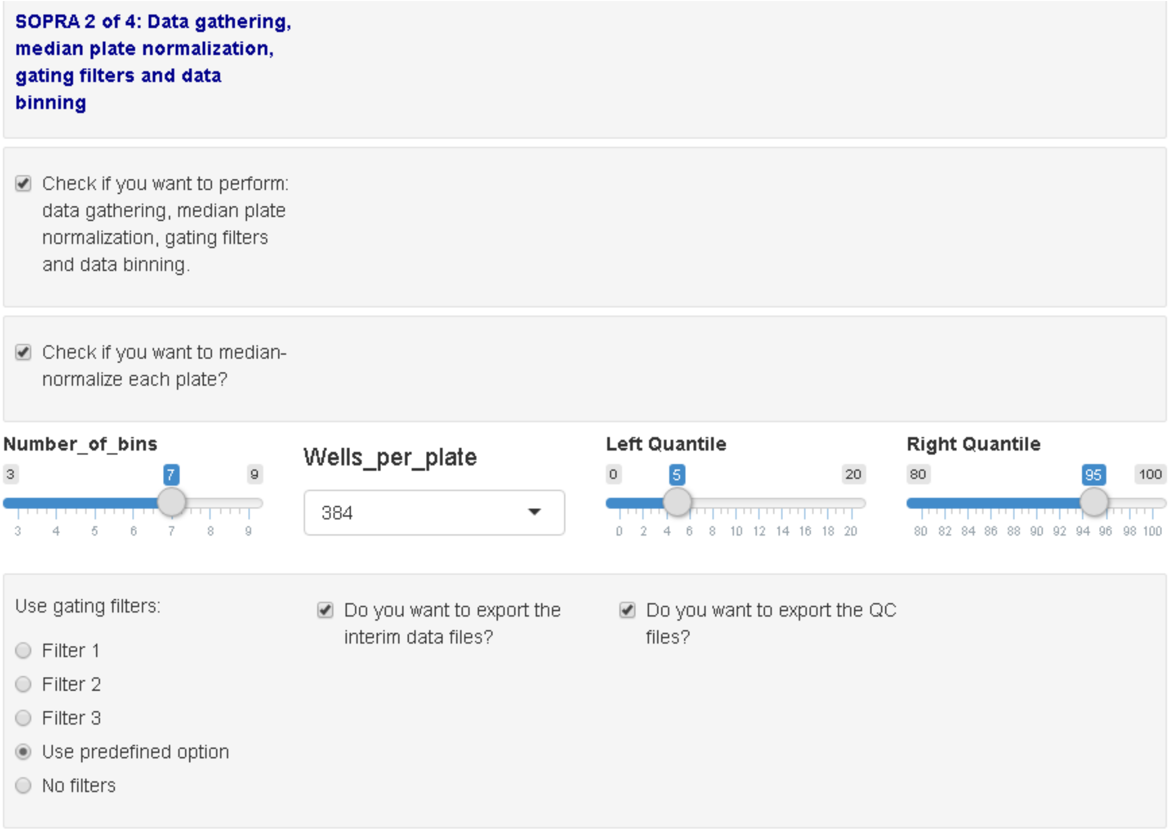


Fig. 9  Menu for SOPRA 2 of 4

#### 4.6 SOPRA 3 of 4 (Find Statistically Significant Profiles- Fig. 10)

As described above, the normalization of the chosen cell population feature leads to density profiles (histograms) for the sample wells relative to the defined control wells. The more distinct a normalized histogram is from a horizontal line, the more significant the change.
The R/Bioconductor package maSigPro is used to perform a regression analysis with a defined false discovery rate, p-value, RSQ-value and degrees of freedom, to identify significantly altered cell population profiles. More information on how the regression analysis is performed can be found in the user's manual of maSigPro at https://www.bioconductor.org/packages/release/bioc/html/maSigPro.html.

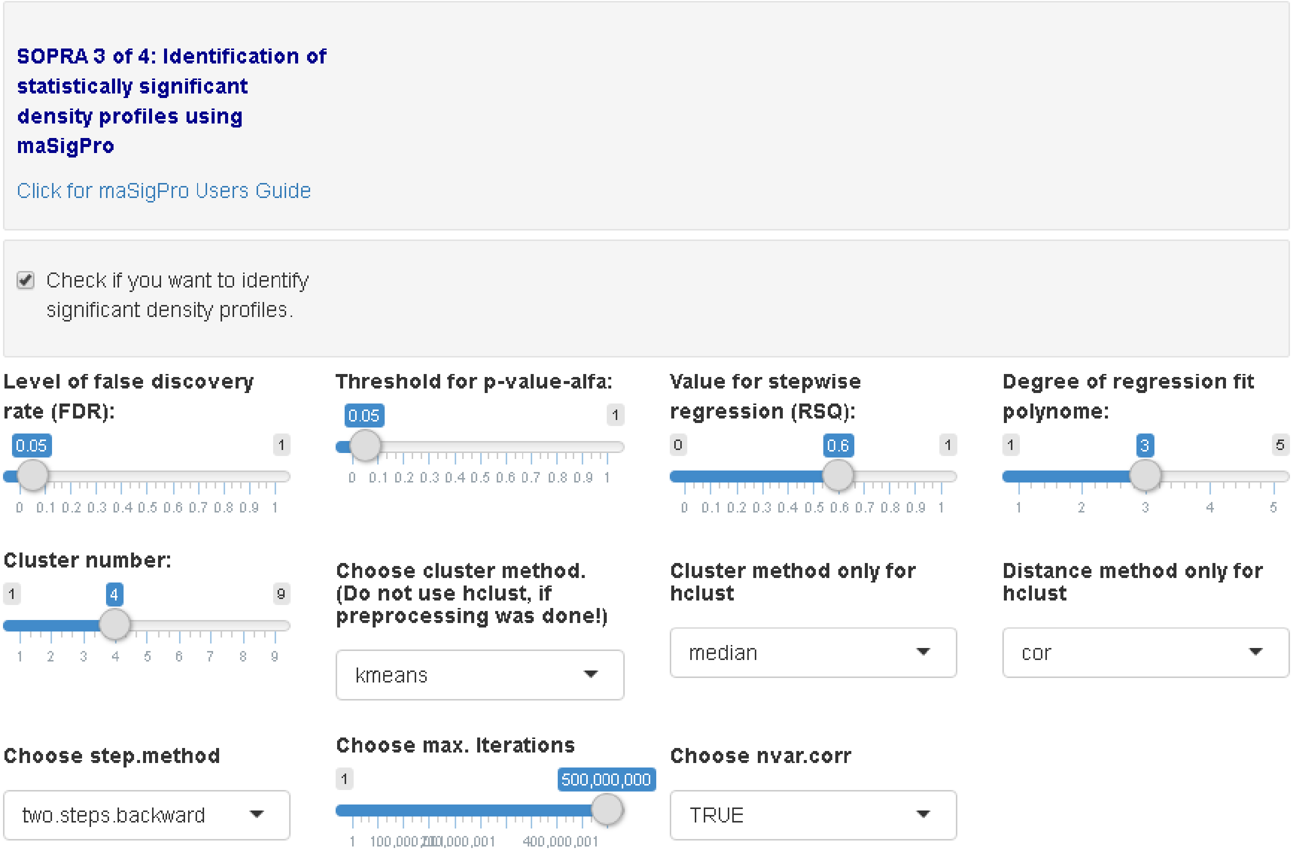


Fig. 10 Menu for SOPRA 3 of 4 (Find statistically significant density profiles)

The result of SOPRA 3 of 4 is exported in the files 'All_p-rsq_values_after_function_tfit_wDegreexQ0.0x_walfa0.0x.txt'.

#### 4.7 SOPRA 3 of 4 (Clustering)

Additionally, the identified statistically significant density profiles can be clustered using a variety of cluster methods (Fig. 11). To read more about the cluster methods, please refer to the maSigPro description.

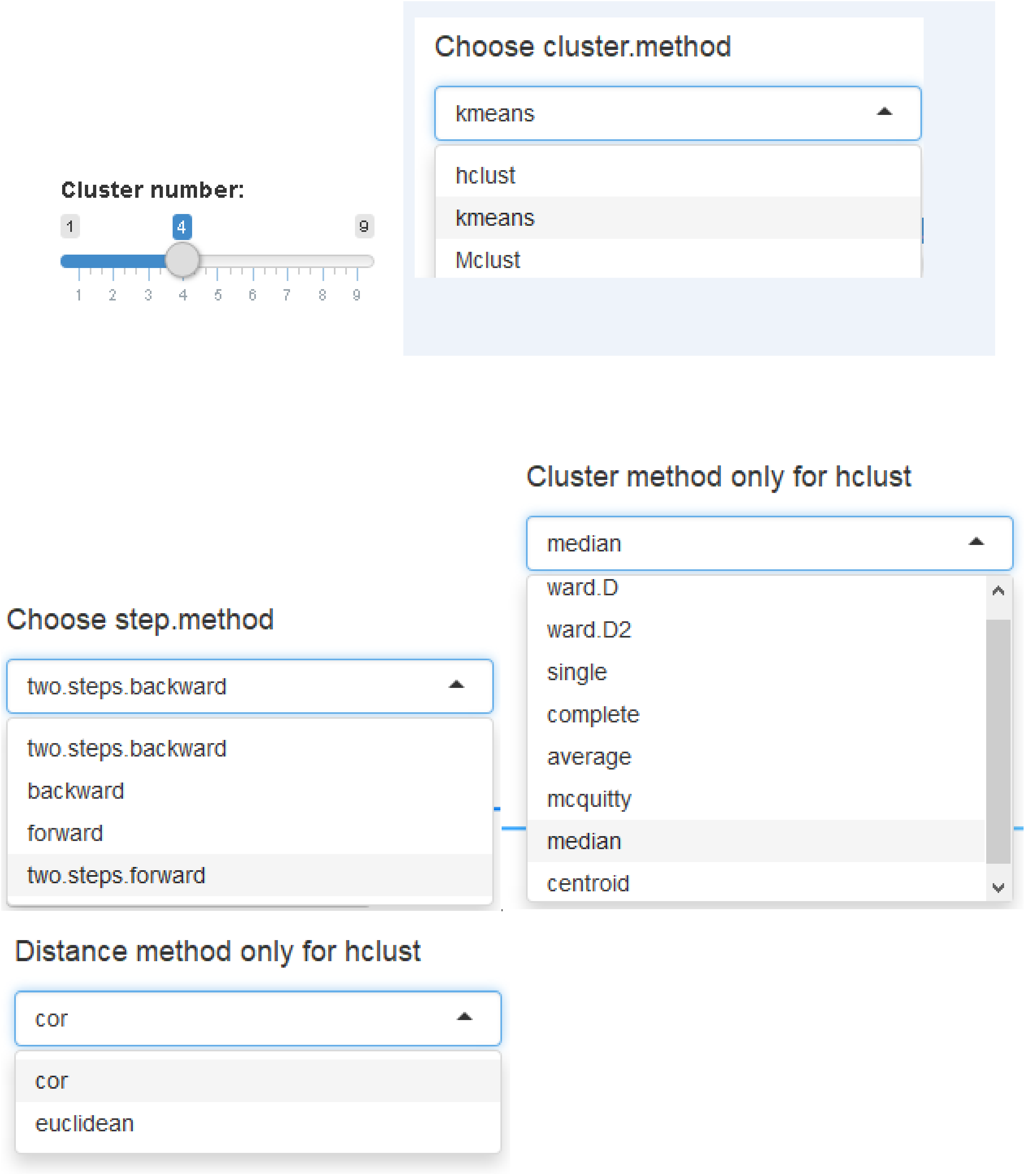


Fig. 11 Selection of cluster parameters for SOPRA 3 of 4

#### 4.8 SOPRA 4 of 4 (Post-processing)

In most siRNA screens multiple siRNA inhibitors are used for the same target gene. Additionally, the same siRNA or chemical compound can be used in several well replicates. The SOPRA workflow uses the following nomenclature to differentiate between multiple siRNAs (or compounds) for the same target gene (or protein) and well replicates: e.g. 'DDX1_siRNA- 1#1#26' for siRNAs and e.g. 'A1#1#88' for chemical compounds.

The first part 'DDX1' or 'A1' is the gene or compound name, the second part 'siRNA-1 'differentiates between different siRNAs used for the same gene. The later part '#1#26' determines the plate and the well location of the inhibitor used.

The SOPRA 4 of 4 (Post-processing- Fig. 12) offers the opportunity to determine which inhibitor was identified in which cluster with which frequency. SOPRA 4 of 4 is running after clustering directly.

The results are exported into the file

"All_significant_siRNA_in_which_cluster__wd3_Q0.0x_alfa0.0x_rsq0.x_clusterx_Mclusttwo.steps.backwardcor_post_processed.txt"

#### 4.9 Start the SOPRA workflow

Finally, after setting all necessary parameters and options, the SOPRA workflow can be started by clicking the 'Run Analysis!' button - Fig. 12)

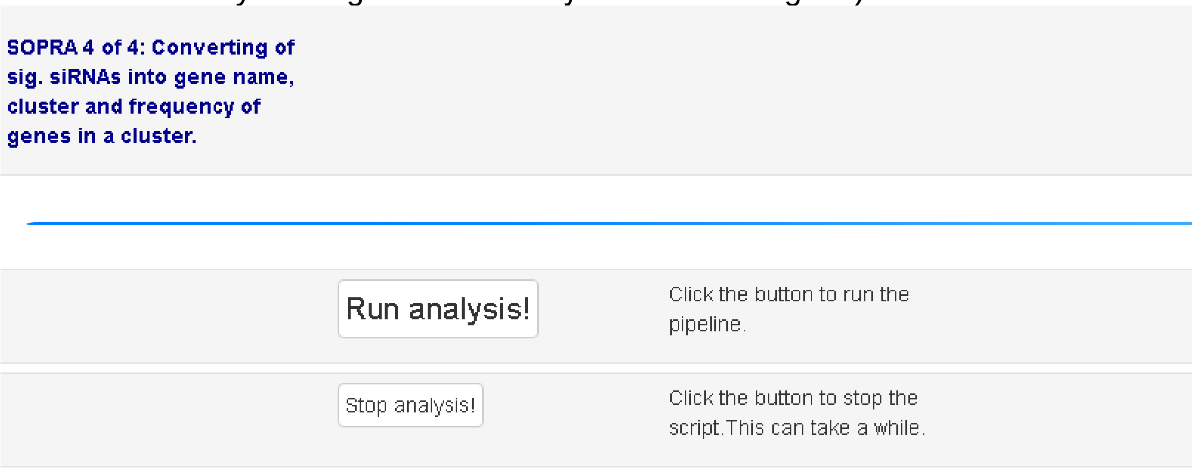


Fig. 12 Post-Processing (SOPRA 4 of 4) and run analysis

After the calculation is finished you can change settings and run the pipeline again. Please note that resulting data files might be overwritten if you use the same output folder. By setting or unsetting the checkboxes one can run the analysis again.

SOPRA 3 of 4 can be rerun independently from SOPRA 1 of 4 and SOPRA 2 of 4 if checkboxes of these steps are unchecked.

The progress of the SOPRA workflow (Fig. 13) can be monitored either by the interactive feedback in the SOPRA window or in the R-Studio console.

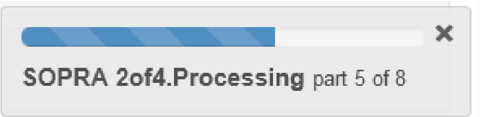


Fig. 13 Interactive feedback in the SOPRA window with progress bar.

If the analysis is finished (Fig. 14) a window pops-up with some explanation how to rerun the workflow under changed conditions or parameters.

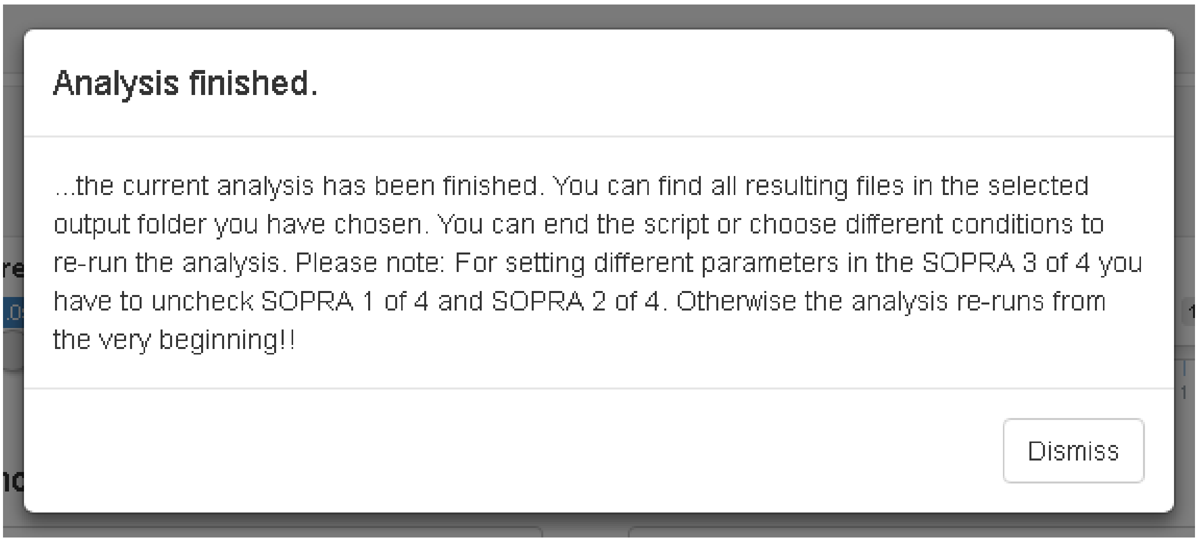


Fig. 14 Analysis finished and some explanations given for re-running the analysis

### 5 Results

#### 5.1 Result of SOPRA 1 of 4 (Preprocessing)

SOPRA 1 of 4 (Preprocessing) flags all objects for wells that are indicated in the ‘ScreenLog.txt’ with ‘NA’. The preprocessed files are saved in the subfolder ’Preprocessing_Results’ in the dedicate output folder.

#### 5.2 Result of SOPRA 2 of 4 (Data Gathering and Normalization)

SOPRA 2 of 4 (Data Gathering and Normalization) results in log2 normalized frequency distributions (histograms) for all samples (and controls). The data is exported in the file ‘AllDataTable.txt’ including all annotations. The file

‘AllDataTable.txt’ contains the processed data, with the plates arranged vertically and the replicates arranged horizontally. The file contains the RNA.ID, plate, replicate and well number, the processing condition (‘TRUE’ or ‘FALSE’) and the binning data followed by the well content and gene symbol.

Optionally, data generated during the calculation process can be exported. The files ‘Data_rawdata_PlateXReplicatey.txt’ contain the raw data values for each object for the feature of interest, while the files ‘Data_nzdata_PlateXReplicateY.txt’ contain the ‘plate-wise’ normalized values for each object for the feature of interest.
For the binning itself three files can optionally be exported, the file ‘Absolute_frequencies_of_PlateXReplicateY.txt’ containing the counts per bin, the file ‘Relative_frequencies_of_PlateXReplicateY.txt’ containing the relative values per bin and the file ‘Binwisenz_frequencies of_PlateXReplicateY.txt’ containing the normalized control values for each bin.

If the analysis is finished the results are depicted via the shiny user interface by clicking on the corresponding tabs (Fig. 15 - Fig. 21).

<< Click on the tabs to see the results >>

Exp.design | QC plot | clustered profiles of sign. siRNAs | sign. siRNAs with cluster | sign. genes with cluster and frequency | Genecount

| Time | Replicate | Bin |
|---|---|---|
| 46 | 1 | 1 |
| 46 | 1 | 1 |
| 46 | 1 | 1 |
| 46 | 1 | 1 |
| 72 | 2 | 1 |
| 72 | 2 | 1 |
| 72 | 2 | 1 |
| 72 | 2 | 1 |
| 98 | 3 | 1 |
| 98 | 3 | 1 |
| 98 | 3 | 1 |
| 98 | 3 | 1 |
| 124 | 4 | 1 |
| 124 | 4 | 1 |
| 124 | 4 | 1 |
| 124 | 4 | 1 |
| 150 | 5 | 1 |
| 150 | 5 | 1 |
| 150 | 5 | 1 |
| 150 | 5 | 1 |
| 176 | 6 | 1 |
| 176 | 6 | 1 |
| 176 | 6 | 1 |
| 176 | 6 | 1 |
| 202 | 7 | 1 |
| 202 | 7 | 1 |
| 202 | 7 | 1 |
| 202 | 7 | 1 |

Fig. 15 Experimental design table necessary for single series time course experiments in maSigPro

Furthermore, quality control plot (QC boxplot) can be exported containing boxplots for all plates after normalization, as well as the distribution of selected cell feature per plate.

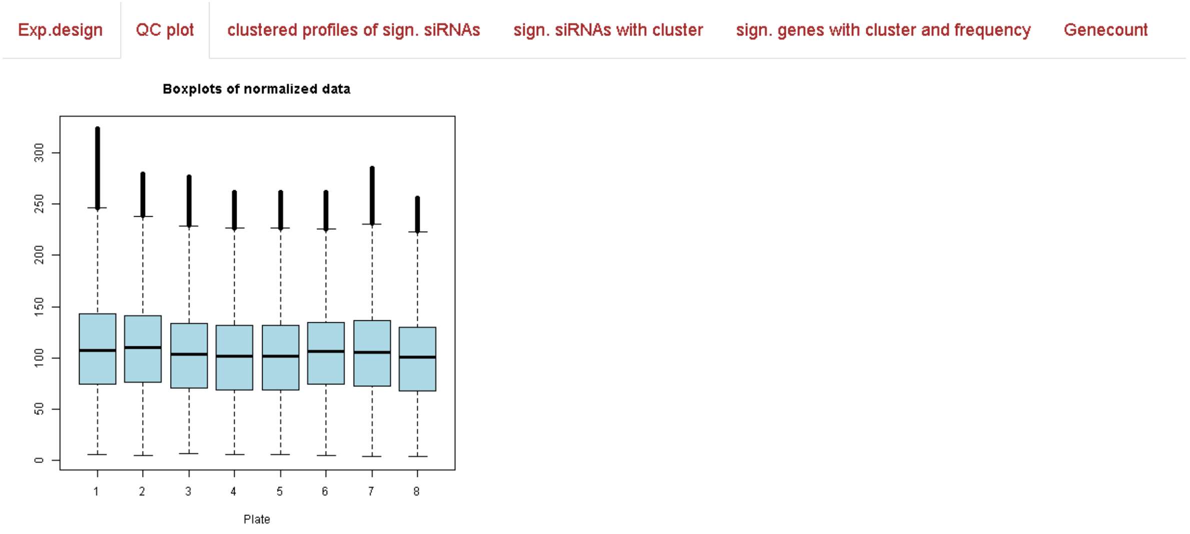


Fig. 16 QC boxplot. It shows the number of plates that were processed with the distribution of selected feature per plate after normalization.

#### 5.3 Result of SOPRA 3 of 4 (Find Statistically Significant Profiles)

For SOPRA 3 of 4 the file 'All_prsq_values_after_function_tfit_wDegreeXQY_walfaZ.txt' is exported into the output folder.This file contains all calculated p-values and rsq-values for each processed probe.

| | p-value | R-squared |
|---|---|---|
| DDX1_siRNA-1#1#26 | 6.25E-11 | 0.873674209 |
| DDX1_siRNA-1#1#27 | 5.56E-08 | 0.776545053 |
| HNRPM_siRNA-15#1#28 | 3.84E-06 | 0.68043179 |
| HNRPM_siRNA-15#1#29 | 1.65E-05 | 0.638235816 |
| PSMC5_siRNA-29#1#30 | 0.001282625 | 0.333834073 |
| PSMC5_siRNA-29#1#31 | 4.61E-08 | 0.689372425 |
| PHB_siRNA-43#1#32 | 0.006574113 | 0.33099868 |
| PHB_siRNA-43#1#33 | 0.008269675 | 0.318604791 |
| BAG1_siRNA-57#1#34 | 4.45E-06 | 0.561593992 |
| BAG1_siRNA-57#1#35 | 4.43E-08 | 0.690277432 |
| BCL10_siRNA-71#1#36 | 0.000547369 | 0.451644245 |
| BCL10_siRNA-71#1#37 | 0.008062588 | 0.31998584 |
| Allstars#1#38 | 0.000770622 | 0.357990056 |
| A1#1#40 | 2.14E-08 | 0.756513428 |
| A2#1#41 | 1.63E-22 | 0.986381108 |
| N1#1#42 | 6.50E-16 | 0.95160789 |
| N2#1#43 | 2.66E-14 | 0.934009892 |
| A3#1#44 | 2.01E-14 | 0.935551082 |
| A4#1#45 | 1.01E-14 | 0.939138851 |
| N3#1#46 | 4.54E-07 | 0.689147363 |
| N4#1#47 | 1.68E-08 | 0.797926353 |

Fig. 17 Result file for SOPRA 3 of 4 with siRNA, p-value and R-squared value

#### 5.4 Result for SOPRA 3 of 4 (Clustering)

Depending on the user's choice, clustering is performed and the results are exported as file "All_significant_siRNA_in_which_cluster__wd3_Q0.05_alfa0.05_rsq0.6_cluster4_hclusttwo.steps.backwardcorgray.txt" and as a png file 'Clustered_Sign_Samples_w….png'. The file name contains hints with which parameters maSigPro was carried out. The clustered profiles of significant siRNAs are depicted (Fig. 18).

Exp.design QC plot clustered profiles of sign. siRNAs sign. siRNAs with cluster sign. genes with cluster and frequency Genecount

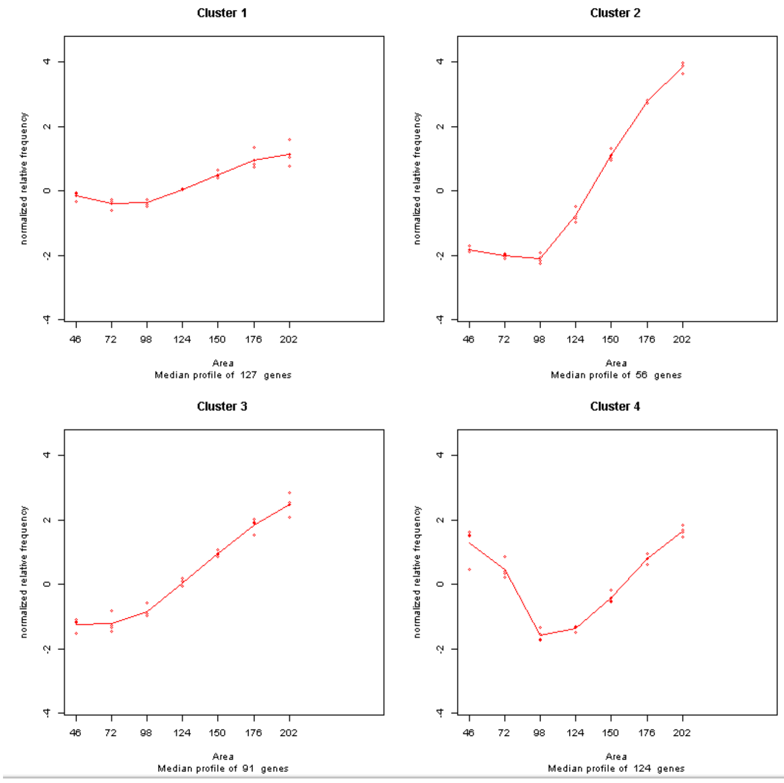


Fig. 18 Statistically significant, normalized frequency density profiles after clustering. Each replicate per plate is vertically depicted as a dot over a binning interval , i.e. here 4 replicates per plate were processed. The x-axis is labeled with the selected cellular feature given in the PlateList.txt

Exp.design QC plot clustered profiles of sign. siRNAs sign. siRNAs with cluster sign. genes with cluster and frequency Genecount

| siRNA | Cluster |
|---|---|
| DDX1_siRNA-1#1#26 | 2 |
| DDX1_siRNA-1#1#27 | 2 |
| HNRPM_siRNA-15#1#28 | 2 |
| HNRPM_siRNA-15#1#29 | 2 |
| PSMC5_siRNA-29#1#31 | 2 |
| BAG1_siRNA-57#1#35 | 2 |
| A1#1#40 | 4 |
| A2#1#41 | 4 |
| N1#1#42 | 3 |
| N2#1#43 | 3 |
| A3#1#44 | 1 |
| A4#1#45 | 1 |
| N3#1#46 | 3 |
| N4#1#47 | 3 |
| BARD1_siRNA-16#1#52 | 2 |
| BARD1_siRNA-16#1#53 | 2 |
| PSMC4_siRNA-30#1#54 | 2 |
| PSMC4_siRNA-30#1#55 | 2 |
| PSMA3_siRNA-44#1#56 | 1 |
| PSMA3_siRNA-44#1#57 | 1 |
| BCL2L11_siRNA-72#1#61 | 2 |
| A1#1#64 | 4 |
| A2#1#65 | 4 |

Fig. 19 Part of statistically significant siRNAs with its cluster membership.

#### 5.5 Result for SOPRA 4 of 4 (Post-processing)

SOPRA 4 of 4 (Post-processing) generates the file 'All_significant_siRNA_in_which_cluster__post_processed.txt' with gene, cluster membership of gene and frequency of gene in the corresponding cluster (Fig. 20).

Exp.design | QC plot | clustered profiles of sign. siRNAs | sign. siRNAs with cluster | sign. genes with cluster and frequency | Genecount

| Gene | Cluster | Frequency |
|---|---|---|
| A1 | 4 | 28 |
| A2 | 4 | 28 |
| N1 | 3 | 28 |
| N2 | 3 | 28 |
| A3 | 1 | 28 |
| A4 | 1 | 28 |
| N4 | 3 | 28 |
| N3 | 3 | 25 |
| PSMA3 | 1 | 4 |
| CSK | 2 | 4 |
| MYC | 2 | 4 |
| RPLP0 | 3 | 4 |
| BET1L | 2 | 4 |
| SF3A1 | 1 | 4 |
| BCL2L1 | 2 | 4 |
| BCL2L11 | 2 | 3 |
| BIRC5 | 3 | 3 |
| BCL2 | 2 | 3 |
| APAF1 | 2 | 3 |

Fig. 20 Result of post-processing showing the gene name, cluster and frequency of gene in the corresponding cluster

Exp.design | QC plot | clustered profiles of sign. siRNAs | sign. siRNAs with cluster | sign. genes with cluster and frequency | Genecount

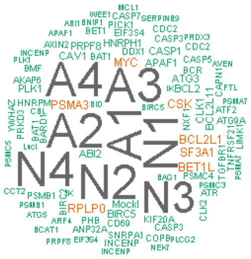


Fig. 21 GeneCount (Wordcloud) shows greater prominence to genes that appear more frequently.

### 6 Software

The software has been implemented in R (https://www.r-project.org) with corresponding R-packages and using RStudio (https://www.rstudio.com) as integrated development environment (IDE).

Software License:
This program is free software: you can redistribute it and/or modify it under the terms of the GNU General Public License as published by the Free Software Foundation, either version 3 of the License, or (at your option) any later version. This program is distributed in the hope that it will be useful, but WITHOUT ANY WARRANTY; without even the implied warranty of MERCHANTABILITY or FITNESS FOR A PARTICULAR PURPOSE. See the GNU General
Public License for more details.

If SOPRA workflow is used to generate data, please cite:

"Single Object Profiles Regression Analysis (SOPRA): A novel method for analyzing high content cell-based screens" Rajendra Kumar Gurumurthy, Klaus-Peter Pleissner, Cindrilla Chumduri, Thomas F. Meyer, Andre P. Maeurer
